# Broken time reversal symmetry in visual motion detection

**DOI:** 10.1101/2024.06.08.598068

**Authors:** Nathan Wu, Baohua Zhou, Margarida Agrochao, Damon A. Clark

## Abstract

Our intuition suggests that when a movie is played in reverse, our perception of motion in the reversed movie will be perfectly inverted compared to the original. This intuition is also reflected in many classical theoretical and practical models of motion detection. However, here we demonstrate that this symmetry of motion perception upon time reversal is often broken in real visual systems. In this work, we designed a set of visual stimuli to investigate how stimulus symmetries affect time reversal symmetry breaking in the fruit fly *Drosophila*’s well-studied optomotor rotation behavior. We discovered a suite of new stimuli with a wide variety of different properties that can lead to broken time reversal symmetries in fly behavioral responses. We then trained neural network models to predict the velocity of scenes with both natural and artificial contrast distributions. Training with naturalistic contrast distributions yielded models that break time reversal symmetry, even when the training data was time reversal symmetric. We show analytically and numerically that the breaking of time reversal symmetry in the model responses can arise from contrast asymmetry in the training data, but can also arise from other features of the contrast distribution. Furthermore, shallower neural network models can exhibit stronger symmetry breaking than deeper ones, suggesting that less flexible neural networks promote some forms of time reversal symmetry breaking. Overall, these results reveal a surprising feature of biological motion detectors and suggest that it could arise from constrained optimization in natural environments.

**Significance:** In neuroscience, symmetries can tell us about the computations being performed by a circuit. In vision, for instance, one might expect that when a movie is played backward, one’s motion percepts should all be reversed. Exact perceptual reversal would indicate a time reversal symmetry, but surprisingly, real visual systems break this symmetry. In this research, we designed visual stimuli to probe different symmetries in motion detection and identify features that lead to symmetry breaking in motion percepts. We discovered that symmetry breaking in motion detection depends strongly on both the detector’s architecture and how it is optimized. Interestingly, we find analytically and in simulations that time reversal symmetries are broken in systems optimized to perform with natural inputs.

## Introduction

Imagine a movie of a car driving (**Fig. 1A**). The car travels into frame from the left, drives along a road, and continues out of frame on the right. Now, imagine the same video, but played in reverse. The car reverses from right to left. The car, originally moving rightwards, has reversed direction and is now moving leftwards. Intuitively, when we reverse time, it seems that our motion percepts should reverse as well (**Fig. 1B**). It is not just our intuition: if we define velocity as displacement over time, playing the movie backward in fact results in equal and opposite velocities everywhere.

**Figure 1.**
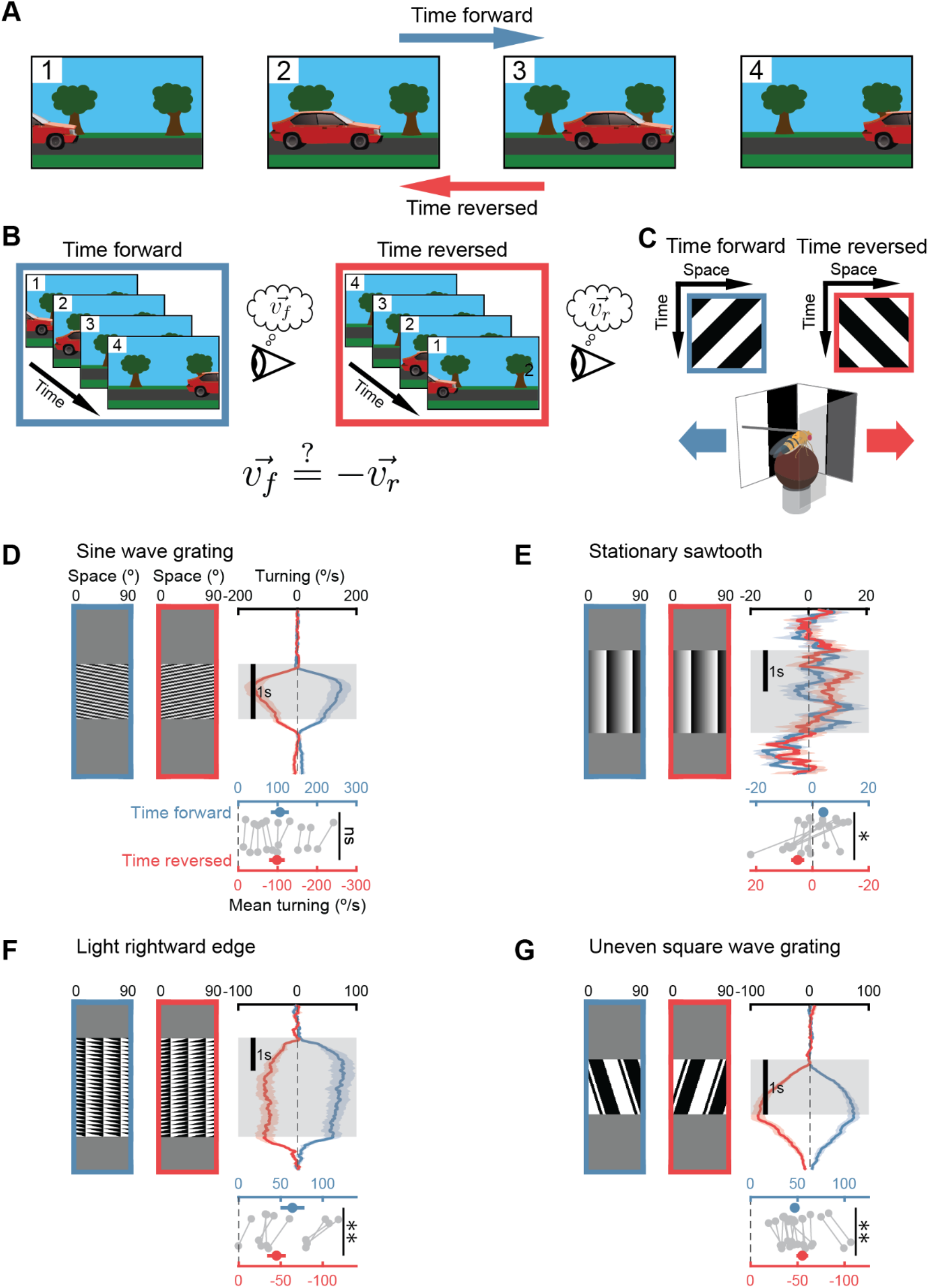
Fly behavioral responses to some stimuli break time reversal symmetry. A. A movie of a car driving forward and rightward becomes a car driving backward and leftward when played in reverse. B. Under time reversal symmetry, the magnitude and direction of motion in a movie is equal and opposite to that when it is reversed in time. Time reversal symmetry of the motion percept is broken when perceived motion vectors are not equal and opposite. C. Motion perception in *Drosophila* was measured with optomotor turning. Tethered flies were placed above air-supported balls while time forward and time reversed stimuli were projected onto panoramic displays. Fly turning was measured by rotation of the ball. D. Responses to drifting sine wave gratings played time forward and time reversed (*left*). Time traces of mean and SEM fly response to each (*top right*). Time traces were averaged to obtain the mean response during the stimulus presentation (*bottom*). Time reversal symmetry breaking is tested by whether the response to the forward stimulus is significantly different from the opposite of the response to the reversed stimulus. N = 9 flies. E. As in (D), but for a stationary contrast sawtooth (21), which is time reversal symmetric. N = 8 flies. F. As in (D), but for local light edges tessellated across space and time. N = 8 flies. G. As in (D), but for square wave grating with bars of unequal widths. N = 8 flies. P<0.05, ^**^ P<0.01 by Student t-test.

Indeed, classical models for biological motion detection match our intuition and produce symmetric responses upon time reversal. The Hassenstein-Reichardt correlator model and the motion energy model, for instance, both estimate the direction and speed of visual motion by computing and averaging pairwise correlations in light intensity over space and time (1, 2). When a stimulus is played in reverse to these models, their output is exactly inverted from when that stimulus is played forward in time (**Appendix 1**). Other models that are optimized to estimate visual motion have similar properties. In particular, a Bayesian model for motion estimation (3) gives a limiting solution that looks like a gradient detector and reverses under stimulus time reversal (**Appendix 1**). One of the most widespread machine-vision techniques for estimating visual motion, the Lucas-Kanade method (4), computes an estimate for local velocities with a method that gives an opposite result when input movies are reversed in time (**Appendix 1**). Thus, a formidable suite of visual motion estimation models, in addition to our intuition, obey what we will call response time reversal symmetry, since reversing time in the stimulus reverses the response, measured by a percept or model output.

However, these models and our intuition stand in contrast to motion perception in real biological circuits, in which responses frequently break time reversal symmetry (see **Figs. 1, S1, S2** for examples).

Symmetries and broken symmetries play an important role in understanding the world and the mathematics used to describe it. In physics, symmetries and invariances play an outsized role in understanding the equations that govern physical phenomena (5, 6). In neuroscience, interesting neural processing properties are often the result of maintaining or breaking symmetries. In visual processing in particular, symmetries and asymmetries of the natural world end up reflected in neural computations. For instance, the translation invariance of visual stimuli across the visual field is reflected in the tiling of feature detectors in vertebrates and invertebrates across visual space (7). In orientation detectors in mammalian cortex, rotational symmetry is broken, so that there are more cells aligned with vertical and horizontal orientations, in agreement with natural scene statistics (8). Also in mammalian cortex, space reversal symmetry, or left-right symmetry, is broken during retinal development to favor front-to-back visual motion that coincides with optic flow due to forward movement (9). Light-dark symmetry is broken in natural scenes, and this is reflected in the number of ON and OFF channels in retina (10) as well as in algorithms for motion detection in flies and vertebrates (11, 12). Moreover, in artificial neural networks, symmetries of the data and the network play critical roles in understanding how machine learning architectures work (13).

Time reversal symmetry has been underappreciated in neuroscience, even though it is critical to understanding dynamical systems like neural circuits (14, 15). Here, we investigate time reversal symmetry breaking in visual motion detection in the fruit fly *Drosophila*. Our goal is to understand the conditions under which a motion percept does not simply reverse when a stimulus is played backward in time. In this study, we first evaluated time reversal symmetry breaking in responses to previously used visual stimuli. We then developed novel methods to allow us to investigate the properties of stimuli that break time reversal symmetry in motion percepts. Using these methods, we discovered a host of new stimuli that break time reversal symmetry in fly motion percepts. To better understand this symmetry breaking, we optimized a suite of simple motion detection models to investigate what properties of those models and their training data result in breaking time reversal symmetry in the model response. We then derived analytically a necessary condition for time reversal symmetry breaking in motion detection. Overall, we have found that motion detection mixes the symmetries in natural scenes; algorithms optimized to detect motion in scenes with non-Gaussian contrasts can break time reversal symmetry, even when time reversal symmetry is maintained in the training data. Our results show how optimization can result in unintuitive regimes of symmetry breaking in neural networks.

## Results

### Certain stimuli elicit behavioral responses that break time reversal symmetry

We have found a variety of visual stimuli that break response time reversal symmetry (**Fig. 1**). To examine directional percepts, we tethered *Drosophila* vinegar flies above an air-suspended ball, surrounded by a panoramic display (**Fig. 1C**) (16). This behavioral assay takes advantage of the optomotor turning response, in which flies and other insects turn in the direction of perceived motion presented across the visual field (1). Some commonly used stimuli, like drifting sinusoidal contrast gratings, do obey time reversal symmetry (**Fig. 1D**). That is, when the drifting grating is played in reverse, the turning response exactly reverses, in line with intuition and with the models discussed earlier (see **Methods, Appendix 1**).

One simple stimulus that elicits percepts that break time reversal symmetry is a stationary sawtooth in contrast (**Fig. 1E**), which elicits motion percepts in humans (17), primates (18), cats (19), fish (20), and flies (21). (This sawtooth stimulus and a different version of this stimulus, designed to create strong illusory motion percept in humans, are shown in **Supp. Fig. S1** (22).) Since this stimulus is stationary, it is identical when played forward or backward, so any net turning response to this stimulus will break time reversal symmetry. Indeed, flies turn in response to this stationary stimulus (**Fig. 1E**), and prior work has shown that this response requires visual neurons that detect motion (21). Flies also respond to non-stationary versions of this stimulus in ways that break time reversal symmetry (**Supp. Fig. S2**).

Other stimuli with strong local motion signals can also elicit behaviors that break time reversal symmetry. For instance, a stimulus consisting of local light edges elicits turning with opposite sign when it is reversed in time, but the turning strengths are not equal (**Fig. 1F**), so this also breaks time reversal response symmetry. This response symmetry can also be broken with a rigid rotation of a pattern. A periodic square wave grating with light and dark bars of unequal widths can elicit responses that are not time reversal symmetric (**Fig. 1G**). This drifting stimulus, similarly to the local light edge stimulus, elicits opposite but unequal turning upon time reversal.

Additional stimuli, including ones that are identical when played forward and backward and that include oppositely oriented light and dark edges (23), also lead to behavior that breaks time reversal symmetry (**Fig. S2**).

### Examining the full suite of stimulus symmetries

The stimuli examined so far result in behavior that breaks time reversal symmetry, but it is less clear what properties a stimulus must have to generate responses that break this symmetry. To analyze what stimulus properties are involved, we began by defining three symmetries in our stimuli: time reversal, space reversal, and contrast reversal. A stimulus possesses a specific symmetry if, when that operation is applied, the stimulus is identical, up to a phase shift in space or time (**Fig. 2A**). For instance, a drifting sinusoidal grating (**Fig. 1D, 2A**) shows a contrast symmetry, since when contrast is reversed, the resulting stimulus is identical to the original up to a phase shift. We represent the time reversal operator as Θ, the contrast reversal operator as Γ, and the space reversal operator as χ(see **Methods** for definitions). Once an operator is applied to a stimulus *S*, we represent the altered stimulus with a left subscript: in other words, Γ[*S*] = _Γ_*S*. In this notation, the stimulus *S* is said to be contrast, time, and space reversal symmetric if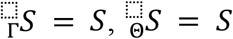, and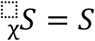, respectively. Time reversal, contrast reversal, and space reversal are commutative and consequently can be noted as sequential left subscripts: for instance,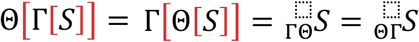. The response, *R*[*S*], represents the integrated turning response of flies to the stimulus *S*. The drifting sinusoid possesses contrast symmetry (Γ symmetry), space-time symmetry (χΘ symmetry), and contrast-space-time symmetry (ΓχΘ symmetry) (**Fig. 2A**).

**Figure 2.**
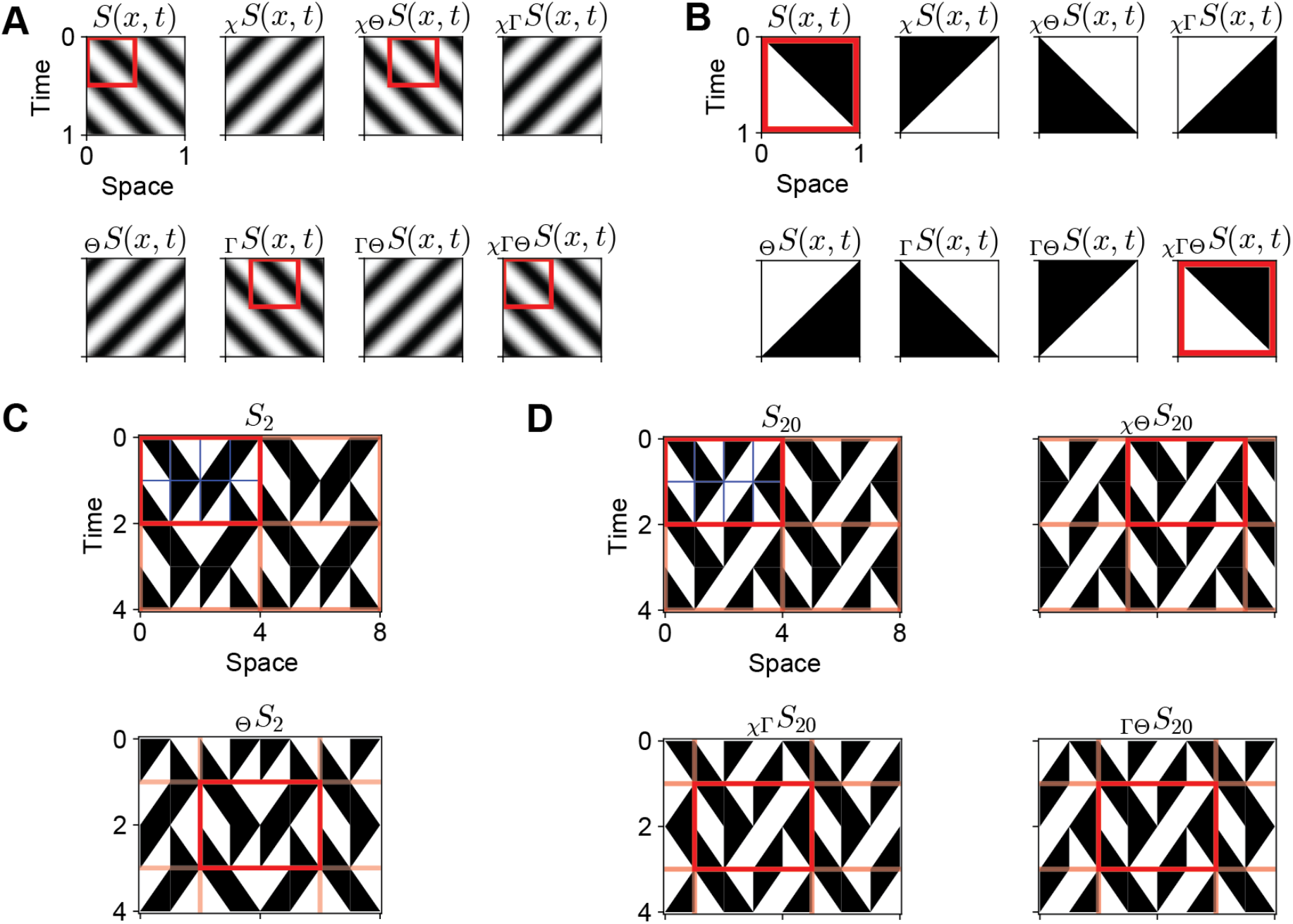
Edge-tile stimuli can be created with different combinations of spatial, temporal, and contrast symmetries. A. All stimuli can be transformed through space reversal, time reversal, or contrast reversal, denoted with operators χ, Θ, and Γ. A stimulus *S* is *O* symmetric if 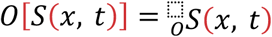 is the same as *S*(*x, t*), allowing for shifts in space and time. A drifting sinusoid grating, shown here, is contrast (Γ) symmetric, space-time (χΘ) symmetric, and contrast-space-time (ΓχΘ) symmetric because the corresponding transformations return the same stimulus tessellated in space and time (boxed in red). B. Edge-tiles consist of single light or dark edges, moving left or right. A light, rightward edge-tile can be transformed into any other single edge-tyle type (light/dark, rightward/leftward) through space reversal, time reversal, or contrast reversal. Each single edge-tile is χΓΘ symmetric. C. Stimuli with different combinations of symmetries can be created by placing arrays of individual edge-tiles into unit cells. *S*_′_ is comprised of a unit cell of 8 edges-tiles in a 4 by 2 grid (outlined in red), which is tessellated over space and time. *S*_′_ is time reversal symmetric because reversing the entire stimulus in time results in a stimulus in which the same unit cell can be found (phase shifted in time and space from its original location and time). (See **Methods** for details on creating these stimuli.) D. Stimuli can possess multiple combinations of symmetries. *S*_′(_ has a unit cell that is 4 by 2 and is χΘ, χΓ, and ΓΘ symmetric, meaning that performing each of those pairs of reversals results in a stimulus possessing the same 4 by 2 unit cell, outlined in red.

We catalogued the symmetries in the initial stimuli we examined (**Figs. 1, S2**) and found that these stimuli covered a relatively small fraction of all possible symmetry combinations (**Table 1**, *top*). Some symmetries, especially space-contrast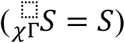, appeared over-represented among those that resulted in behavior that broke time reversal symmetry.

**Table 1.**
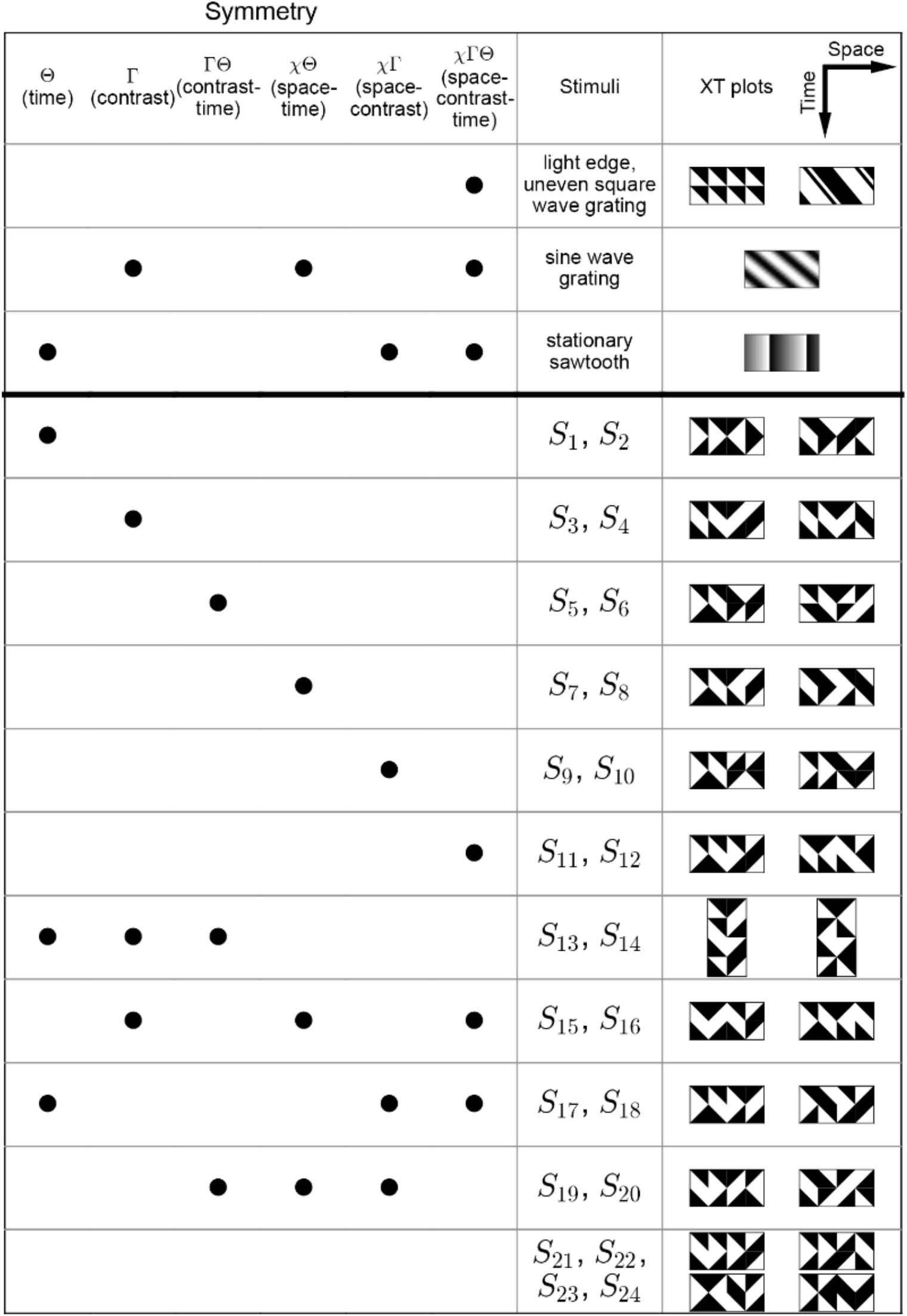
Table of symmetries in stimuli in Figure 1. The upper part of the table enumerates symmetries in our stimuli from Figure 1, while the lower part enumerates those in the edge-tile stimuli. A dot in the column means that the stimulus possesses that symmetry. (See **Methods** for the definition of the symmetries with periodic stimuli.) The different symmetries are Θ (time reversal), Γ (contrast reversal), χ(space reversal), ΓΘ (contrast-time reversal), χΘ (space-time reversal), χΓ (space-contrast reversal), and χΓΘ (space-contrast-time reversal).

| $\Theta$<br>(time) | $\Gamma$<br>(contrast) | $\Gamma\Theta$<br>(contrast-time) | $\chi\Theta$<br>(space-time) | $\chi\Gamma$<br>(space-contrast) | $\chi\Gamma\Theta$<br>(space-contrast-time) | Stimuli | XT plots |
| --- | --- | --- | --- | --- | --- | --- | --- |
|  |  |  |  |  | ● | light edge, uneven square wave grating |  |
|  | ● |  | ● |  | ● | sine wave grating |  |
| ● |  |  |  | ● | ● | stationary sawtooth |  |
| ● | | | | | | $S_1, S_2$ | |
| | ● | | | | | $S_3, S_4$ | |
| | | ● | | | | $S_5, S_6$ | |
| | | | ● | | | $S_7, S_8$ | |
| | | | | ● | | $S_9, S_{10}$ | |
| | | | | | ● | $S_{11}, S_{12}$ | |
| ● | ● | ● | | | | $S_{13}, S_{14}$ | |
| | ● | | ● | | ● | $S_{15}, S_{16}$ | |
| ● | | | | ● | ● | $S_{17}, S_{18}$ | |
| | | ● | ● | ● | | $S_{19}, S_{20}$ | |
| | | | | | | $S_{21}, S_{22}, S_{23}, S_{24}$ | |

To examine how stimulus symmetries affect the response symmetries, we set out to design a suite of stimuli that could fill in all possible combinations of symmetries in **Table 1**. To do that, we established a new method to design edge-tile stimuli (see **Methods**). These stimuli use edge-tiles that contain either light or dark edges, moving either to the right or to the left (**Fig. 2B**). After arranging groups of 8 individual edge-tiles into rectangular unit cells, we could tesselate the unit cells in space and time (**Fig. 2C, D**). Since there are 4 possible edge-tiles at each location in the unit cell, there are 4^8^ = 65,536 possible unit cells, or patterns, though not all result in unique stimuli after tessellation. To check for different symmetries in stimuli made with each pattern, we compared the stimulus transformed with the symmetry operators to the original stimulus at all possible phase shifts (for most patterns, 4 in space, 2 in time; see **Methods**). If the transformed and original stimuli contain the identical unit cell at some phase offset, then the stimulus possesses that symmetry. For each unit cell, we enumerated whether it has a space reversal symmetry, time reversal symmetry, contrast reversal symmetry, or various combinations of these (**Fig. 2C, D**).

Next, we selected 24 different edge-tile stimulus patterns to test on flies. For each possible combination of symmetries, we chose two patterns (**Table 1**, *bottom*). Four patterns were chosen with no symmetries. Where possible, the specific patterns were selected to minimize contrast discontinuities in time, for instance a dark region that instantaneously becomes light. Also where possible, stimuli for each combination of symmetries were selected to cover cases where stimuli had both non-zero and zero net motion when averaged over time and space.

### Fly responses to certain edge-tile patterns break time reversal symmetry

We presented each of the chosen edge-tile patterns to wildtype *Drosophila melanogaster* flies. Since the patterns have only one spatial dimension, we presented them on screens by replicating the same pattern on all pixel rows vertically (see **Movie 1**). Flies responded to many of these stimuli, turning on average more in one direction than in the other (**Fig. 3A, B**). As we did before, we averaged the turning responses over time and over trials to obtain a single mean response to each stimulus for each fly (**Fig. S3**). We began by examining responses to two example stimuli, *S*_2_ and *S*_20_, which elicited responses that broke time reversal symmetry (**Fig. 3A, B**). We plotted responses to different presentations of the stimuli (**Fig. 3i, ii**). To analyze symmetry breaking in these responses, we defined three metrics to identify time, contrast, and time-contrast reversal symmetry breaking (**Fig. 3iii**). First, the time reversal symmetry metric: a response breaks time reversal symmetry if the response to the stimulus played forward plus the response to it played backward is different from zero, that is, when the metric, 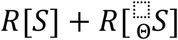, is different from 0. This formulation is the same definition used in our earlier analysis (**Fig. 1**). Second, since one might expect reversing contrast not to change motion percepts, we define the contrast reversal symmetry metric differently: a response breaks contrast reversal symmetry if the response to the stimulus minus the response to the contrast reversed stimulus is different from zero, that is, when the metric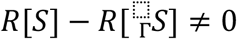. Breaking contrast-time reversal happens when the response to the stimulus plus the response to its contrast and time reversed variant is different from zero, that is, when the metric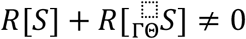. When these metrics were statistically significantly different from zero, we judged the symmetry broken for that stimulus (**Fig. 3iii**). We did not test for space reversal symmetry: when mounting flies, we expect some mild left-right asymmetries in how they are mounted, which we do not want to mistake for true biological asymmetries. Thus, we compute fly responses as half of their turning to the stimulus presented clockwise minus half their turning when it is presented counter-clockwise (see **Methods**). This eliminates potential experimental asymmetry and also prevents us from measuring space reversal symmetry breaking in the response.

**Figure 3.**
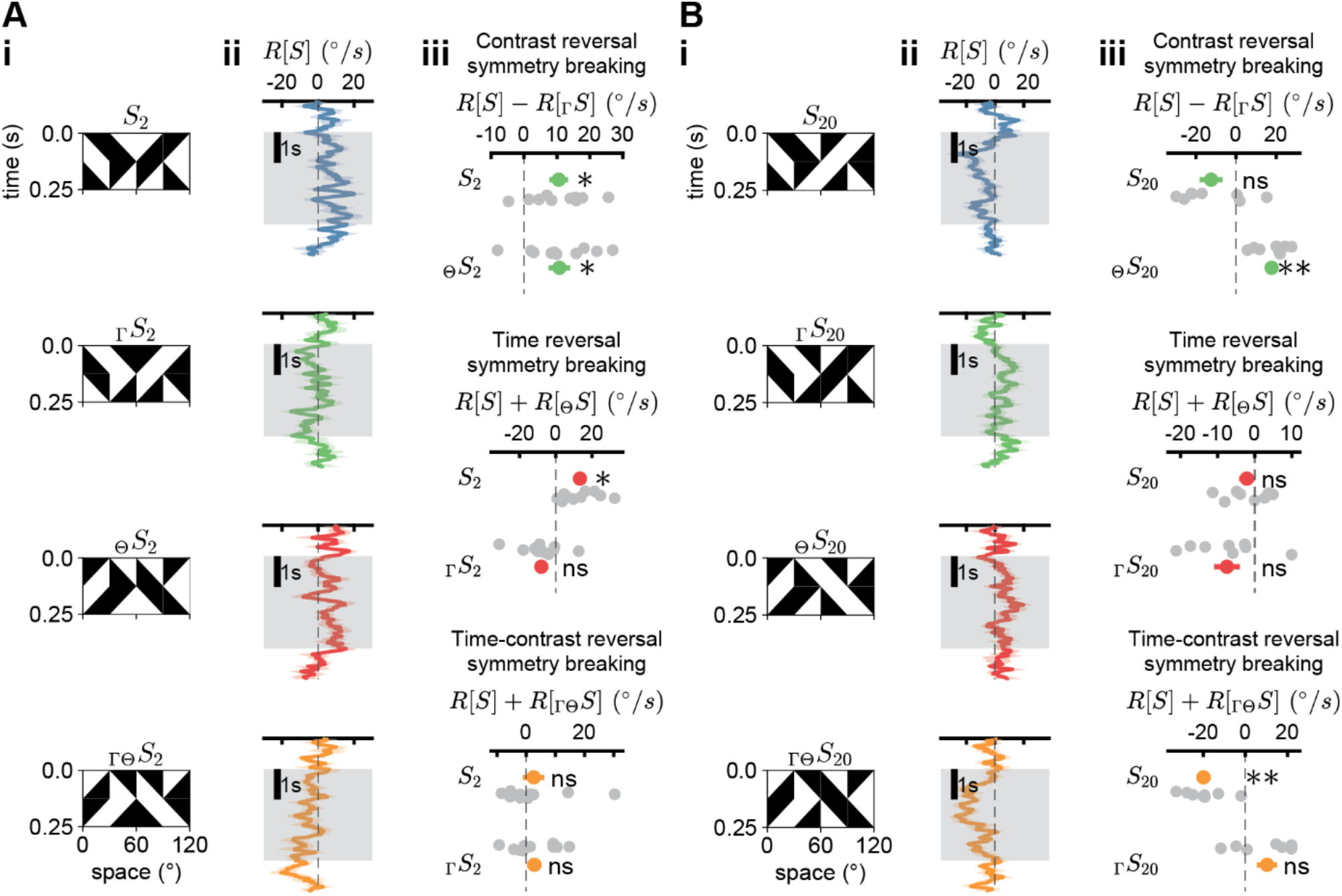
Fly responses to edge-tile stimuli can break time reversal symmetry. Edge-tile stimuli were presented for a total of three seconds, with each individual edge moving at 240°/s. *N* = 8-10 flies for each trace. (^*^ P<0.05, ^**^ P<0.01 by Student t-test, with Holm-Bonferroni correction for 6 comparisons for each stimulus.) A. Fly responses to the stimulus *S*_′_, which is Θ symmetric but generates responses that break time reversal symmetry. (i) The stimulus and its reversals in time, contrast, and time-contrast. (ii) Time traces of mean fly responses to these four transformations of the stimuli. (iii) Contrast symmetry breaking is measured by subtracting the time averaged responses to the original stimulus and the contrast reversed stimulus (green, *top*). Time and time-contrast symmetry breaking is measured by summing time-averaged responses to the original and time reversed or time-contrast reversed stimuli, respectively (red and orange, *bottom*). B. As in (A), but for the stimulus, *S*_′(_, which is χΘ, χΓ, and ΓΘ symmetric.

### Summarizing responses to stimuli with different symmetries

To better understand how specific symmetries in the stimuli influenced responses, we first sorted the stimuli in order of the strength of time reversal symmetry metric and time-contrast reversal symmetry metric (**Fig. 4A,B**). To do this, we plotted the signed symmetry metrics for each stimulus: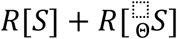 and 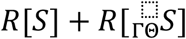. Given asymmetries in the fly’s turning to light and dark edges (**Fig. 1**) (11, 23-26), one might hypothesize that the turning to our edge-tile patterns could be predicted just by examining the number of light and dark edges. However, we did not observe any obvious patterns in the strength of the symmetry breaking as a function of the net number of light and dark edges in the stimuli.

**Figure 4.**
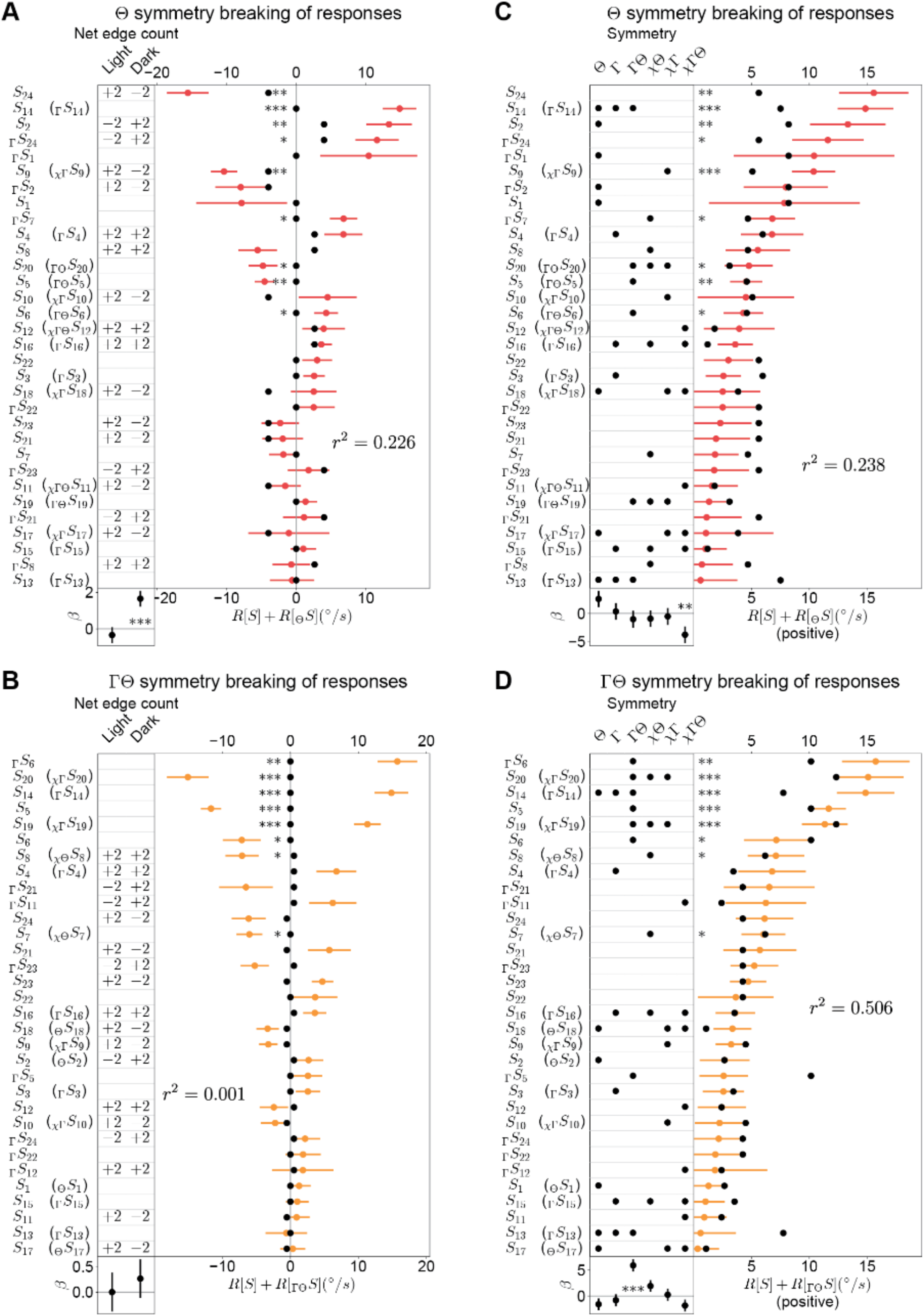
Many stimuli elicit behavior that breaks time reversal symmetry, but stimulus symmetries only modestly predict degree of time reversal symmetry breaking. *N* = 8-11 flies for each measurement. For the data, error bars are 1 SEM; ^*^ P < 0.05, ^**^ P < 0.01, ^***^ P < 0.001 by Student t-test, with Holm-Bonferroni correction for 6 comparisons for each stimulus. For the linear fits, error bars are 1 SEM; ^*^ P < 0.05, ^**^ P < 0.01 ^***^ P < 0.001 by standard confidence interval estimation. A. Counts of net light and dark edges (*left*) and time reversal symmetry metric (*right*). A linear model with no bias term was fit to predict the response from the net light and dark edges. Its predictions are shown as black dots and the fitted weights, *β*, are shown at bottom. The variance accounted for by the model is noted by the *r*^2^ value. Stimuli in parentheses indicate that the stimulus being tested possesses a symmetry (or multiple symmetries) that permit combining response asymmetry metrics from both *S* and 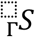 into a single measurement. B. As in (A), but for time-contrast reversal symmetry breaking. C. Symmetries belonging to each stimulus (left) and time reversal symmetry breaking degree (*right*). Compared to (A), all negative time reversal symmetry metrics are flipped in order to make them positive. A linear model with a bias term was fit to predict the degree of symmetry breaking for each stimulus from its set of symmetries. Its predictions are shown as black dots and the fitted weights, *β*, are shown at bottom. The variance accounted for by the model is noted by the *r*^2^ value. As in (C), but for time-contrast reversal symmetry breaking.

To more quantitatively assess the relationship between the time reversal symmetry metric and the count of light and dark edges in the stimuli, we fit linear models to predict the metric by weighting the net light and dark edge count in the edge-tile stimuli. The linear models accounted for 23% and 0.1% of the variance in the time reversal symmetry and time-contrast reversal symmetry metrics. Significant positive weighting was placed on the net dark edges in the case of time reversal symmetry (**Fig. 4A**). No significant weighting was placed on net light or dark edges in the case of time-contrast reversal symmetry (**Fig. 4B**). No constant bias term was fit in the model, as we expect rightward edges of a given polarity to yield the exact opposite percept as leftward edges of the same polarity. Notably, the stimulus *S*_14_ had net 0 light and dark edges but fly responses showed strong symmetry breaking with stimulus time reversal. The six strongest responses breaking time-contrast symmetry all also had net zero light and dark edges. Similarly, many stimuli that broke contrast reversal symmetry in responses also had no net light or dark edges (**Supp. Fig. S4**). Because light and dark edge counts do not predict time reversal symmetry breaking of behavioral responses, the phenomenon is not simply a result of differing strengths of responses to light and dark edges (see also **Supp. Fig. S2** for a non-predictive example).

We next examined how different stimulus symmetries impacted the strength of response symmetry breaking. Since the symmetries in a stimulus have no direction, we flipped all the measured metrics to have a mean positive response (**Fig. 4C, D**). We could then ask how the stimulus symmetries affected the amplitude of the symmetry metrics for both time reversal and time-contrast reversal symmetries. Again, we did not see an obvious pattern in the stimulus symmetries that elicited the strongest response symmetry breaking. We therefore fit linear models to predict the strength of the symmetry breaking from the suite of stimulus symmetries. These models predicted 24% and 51% of the variance in the time reversal and time-contrast reversal behavioral symmetry breaking. The most significant weight was a negative weighting on time-space-contrast symmetry for the response time reversal symmetry breaking (**Fig. 4C**) and a positive weighting on time-contrast symmetry for the response time-contrast reversal symmetry breaking (**Fig. 4D**). This second case shows that flies exhibited net turning to stimuli that were identical when transformed by reversing both time and contrast. This is conceptually similar to flies turning to stimuli that are identical under time reversal (**Figs. 1, S2**). Overall, stimulus symmetries showed some correlations but did not yield definitive contributions of specific properties to breaking time reversal symmetry in fly behavior.

### Optimized neural networks predict stimulus velocity

Stimulus symmetries did not to provide definitive answers about what stimulus features produce responses that break time reversal symmetry, so we opted to take a parallel approach. In this approach, instead of focusing on stimulus properties directly, we worked to understand what factors cause certain motion detectors to break time reversal symmetry. Biological motion detectors have evolved under natural selection, and we wondered how that process of optimization could influence time reversal symmetry breaking in responses. We thus set out to construct neural network models and train them to predict the velocity of a scene. Our goal was not to exactly reproduce the fly’s visual responses, but rather to understand what features of the neural network and training regime result in breaking the response’s time reversal symmetry.

We began by creating a set of synthetic motion stimuli by rigidly translating panoramic images from a database of natural scenes (27) (**Fig. 5A**). For simplicity, we considered only motion in the horizontal direction, mimicking the fly’s rotation in the yaw dimension that we measured in our earlier experiments. Velocity trajectories were drawn from a Gaussian distribution with a correlation half-decay time of 200 ms (**Fig. 5B**). Critically, this dataset is statistically identical when played forward or backward in time, so any time reversal symmetry breaking observed in model responses cannot be the result of broken time reversal symmetry in the training data. For each data sample with a certain velocity trace (**Fig. 5B**), we created a counterpart with spatial coordinates reversed horizontally so that the training dataset had perfect left-right symmetry (**Fig. 5C**). We then trained convolutional neural network models on this motion dataset, so that the models predicted the instantaneous scene velocity based on recent observations of the stimulus contrast over time (**Fig. 5C**). We began with a simple model, which also resembles the motion detectors in the fly eye. This model had two channels that each received inputs from three adjacent points in space, resembling the ON and OFF motion detectors in the fly’s eye (25, 28). This model is similar to prior models used in simulations (29) and task optimization (30). The training converged after ∼300 epochs (**Supp. Fig. S5**) and typical trained models predicted ∼90% of the variance in scene velocity of a hold-out test dataset (**Supp. Fig. S5**). Since these models have no unique optimized solution, we trained 500 models, each with different initializations, to generate a rich group of trained motion detectors to analyze.

**Figure 5.**
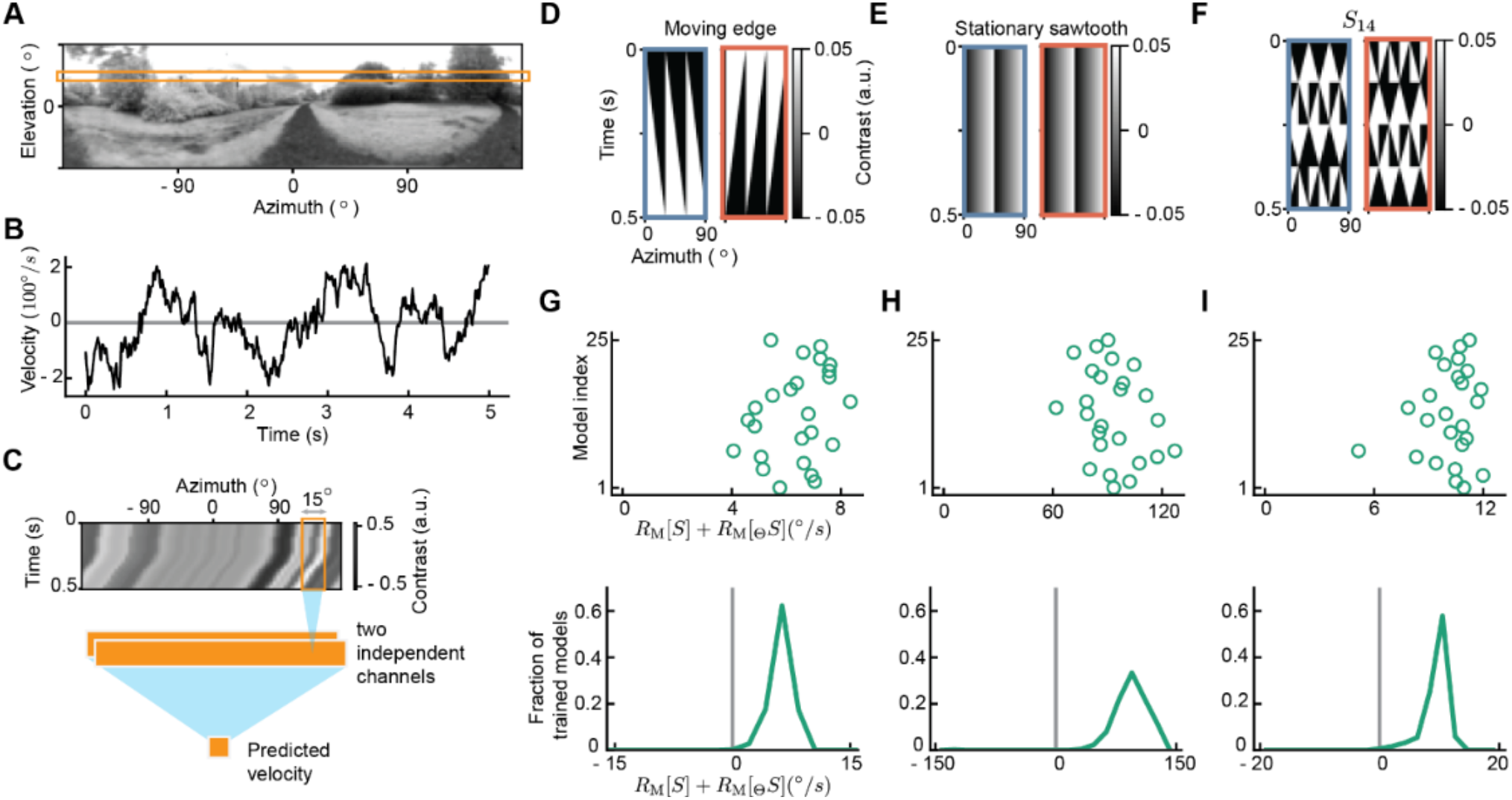
Optimizing an artificial network to detect naturalistic motion yields solutions that break time reversal symmetry. A. An example panoramic natural scene image (27). The elevation spans –48º to 48º and the azimuth spans –180º to 180º. B. An example velocity trajectory with autocorrelation half-decay time of 0.2 s. C. Model architecture. The input to the model is motion stimuli generated by combining a horizontal slice of an image (as in (A)) with 0.5 s of a velocity trajectory (as in (B)). The model output is the predicted velocity at the last time point of the 0.5 s velocity trajectory. There are two independent channels, and each channel spans 3 neighboring inputs in azimuth, 5º apart. (See **Methods** for details.) D. Moving light edge stimulus played forward (*left*) and reversed (*right*) in time. E. Stationary contrast sawtooth stimulus played forward (*left*) and reversed (*right*) in time. Since the stimulus is stationary, the two versions are identical. F. *S*_14_played forward (*left*) and reversed (*right*) in time. Since the stimulus has time reversal symmetry, the two versions are identical up to a phase shift. G. Time reversal symmetry metric for the model response, defined as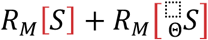, where *R*_*M*_[*S*] is the time averaged response to the stimulus *S*. 25 trained models with different initializations are shown (*top*). Distribution of 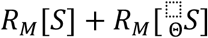 for all trained models evaluated on the moving edges (*bottom*). Only models with testing r-squared values larger than 0.8 are included. H. As in (G), but with the model acting on the stationary sawtooth stimulus. I. As in (G), but with the model acting on *S*_14_.

### Optimized model responses break time reversal symmetry

To investigate time reversal symmetry in the responses of these trained models, we tested them on three synthetic stimuli that had resulted in time reversal symmetry breaking in fly behavior (**Fig. 5D, E, F**): (1) moving light edges and their time reversed counterparts; (2) stationary sawtooth contrast stimuli; and (3) the stimulus *S*_14_, which elicited strong turning responses and possesses time, contrast, and time-contrast reversal symmetry.

The trained models responded more strongly to light edges than to time-reversed light edges (dark edges), indicating that they had broken time reversal symmetry in their responses (**Fig. 5G**). The degree of symmetry breaking here was measured by the time reversal symmetry metric for the model’s response, 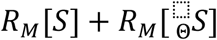, where *R*_*M*_ is the model’s response. This metric would be zero for a response showing time reversal symmetry and non-zero otherwise.

When presented with a stationary sawtooth stimulus or with the stimulus *S*_14_, a model with perfect time reversal symmetry will generate a response of zero, while a model that breaks time reversal symmetry will have a non-zero response. When our trained models were presented with sawtooth stimuli or *S*_14_, they had non-zero responses and time reversal symmetry metrics (**Fig. 5H, I**), indicating the responses broke time reversal symmetry.

For all three stimuli, these trained models broke time reversal symmetry in the same direction as the fly data (**Figs. 1, 4**): the model’s time reversal symmetry metrics, 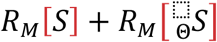, were the same sign as the fly’s metrics, 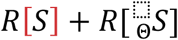 (**Fig. 1**). However, this agreement should not be viewed as significant, since it depends on the stimulus parameters used, such as contrast levels (**Supp. Fig. S5**). Rather, the important result is that these trained models broke time reversal symmetry.

### Time reversal symmetry breaking depends on the contrast distribution in the training data

We wanted to understand which properties of the models and the training data led to breaking time reversal symmetry in the responses. We therefore created synthetic image datasets with contrast distributions different from the natural scenes we used earlier (**Fig. 6A**). In natural scenes, the distribution of the pixel contrast values is right-skewed and highly kurtotic (**Fig. 6A**) (31). This skew has been proposed to explain a suite of fly directional behavioral responses (32).

**Figure 6.**
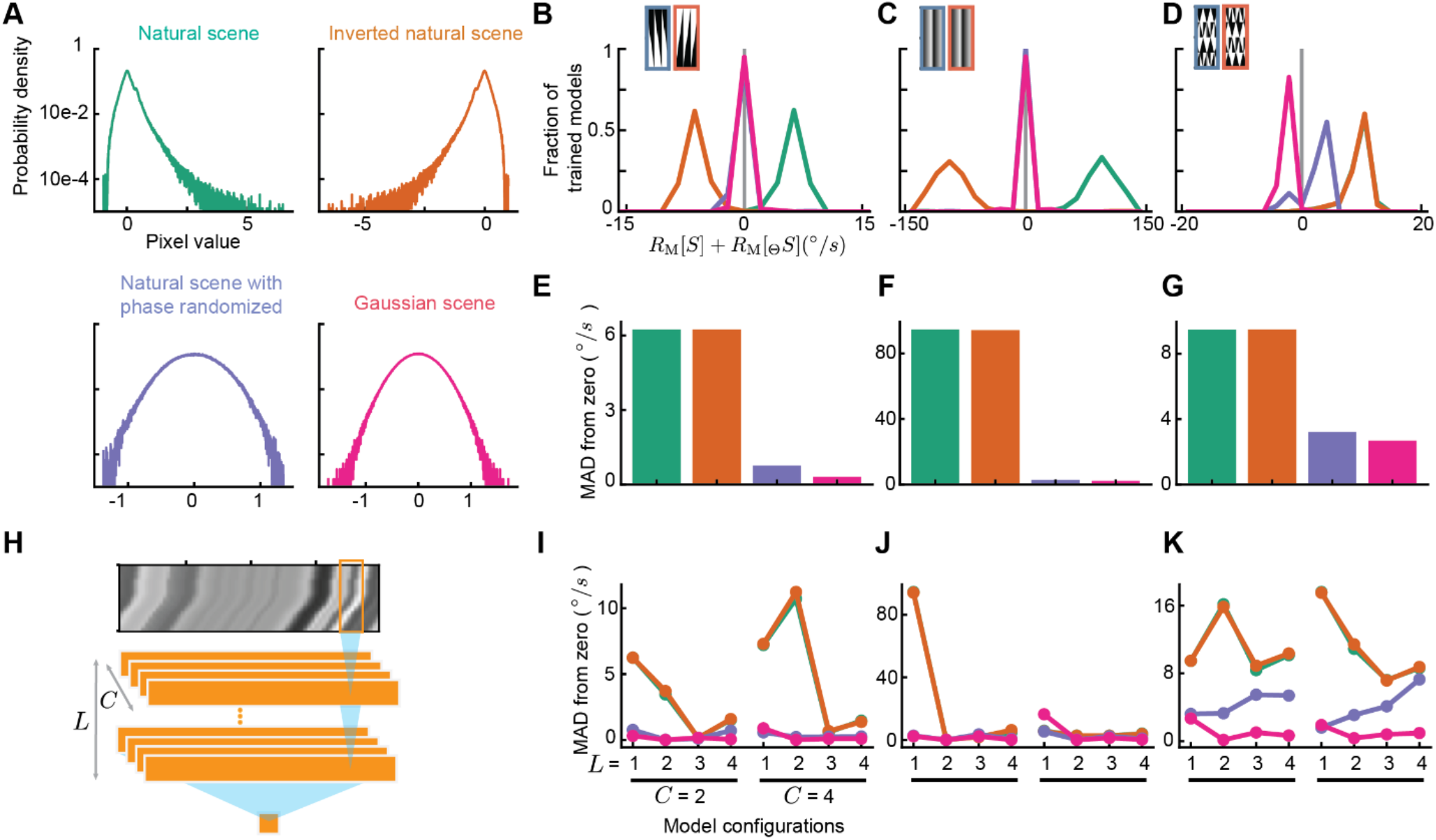
Contrast distributions in training data and shallow architectures drive time reversal symmetry breaking. A. Distributions of the pixel values in the four types of training data: natural scenes, contrast reversed natural scenes, natural scenes with randomized phase, and Gaussian scenes (see **Methods**). B. Distributions of the time reversal symmetry metric,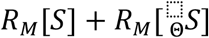, for models trained on the four types of training data in (A). The models are acting on the light edge stimulus. Models are only included if they achieved an r-squared of >0.8 on their test data. C. As in (B), but for the models acting on the stationary sawtooth stimulus. D. As in (B), but for the models acting on *S*_14_. E. Mean absolute deviation (MAD) from zero of the time reversal symmetry metric 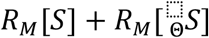 for moving edges. F. As in (E), but for stationary sawtooth stimuli. G. As in (E), but for *S*_14_. H. Model architecture was made variable so that it possessed different numbers of channels *C* and layers *L*. I. Mean absolute deviation (MAD) from zero of the time reversal symmetry metric for moving edges, but for models with different numbers of channels *C* and layers *L*. J. As in (I), but for stationary sawtooth stimuli. K. As in (I), but for *S*_14_.

We therefore decided to manipulate the pixel distribution in the training data to see how it affects response time reversal symmetry breaking. To do this, we created three synthetic training datasets (**Fig. 6A**). The first one simply reversed the sign of the pixel values of the natural scene training set. The second one randomized the phases of Fourier components of the natural scene images, which resulted in images that had symmetric but non-Gaussian pixel value distributions, retaining the spatial correlations of the original images. A third synthetic dataset was created by randomly sampling pixel values from a Gaussian distribution. In the two randomized datasets, the pixel variance was chosen to match the variance in the natural scenes.

Next, we trained our simple model on these three sets of synthetic scenes. All the trained models could perform well on their corresponding test datasets, often predicting 90% of the velocity variance (**Supp. Fig. S6**). To test whether these new training datasets resulted in models whose responses broke time reversal symmetry, we then presented the trained models with the edge stimulus, the stationary sawtooth stimulus, and *S*_14_. When presented with edges or with the sawtooth stimulus, which possess contrast asymmetries, the models trained on the contrast reversed natural scene data broke time reversal symmetry, but in the opposite direction of the models trained on the original natural scene data (**Fig. 6B, C**). For those same stimuli, the models trained with phase randomized and Gaussian scenes had responses that preserved time reversal symmetry (**Fig. 6B, C**). In the case of *S*_14_, which is itself contrast reversal symmetric, the contrast reversed and natural scene trained models gave similar responses, which broke time reversal symmetry (**Fig. 6D**). Interestingly, the models trained on phase randomized and Gaussian scenes also had responses that broke time reversal symmetry, though less strongly than the other two datasets, since the time reversal symmetry metric was closer to 0 (**Fig. 6D**).

We quantified the degree of response symmetry breaking by finding the mean absolute deviation from 0 of the time reversal symmetry metric over all the models trained on each dataset (**Fig. 6E, F, G**). This confirmed the strong symmetry breaking by models trained on natural scene statistics, and the negligible symmetry breaking by models trained on the phase randomized and Gaussian scenes, except in the case of *S*_14_.

Last, we wanted to understand how model architecture influenced the degree of time reversal symmetry breaking. We therefore trained models that had different numbers of channels and layers (**Fig. 6H**). We trained these models on the four datasets and measured the degree of time reversal symmetry by the mean absolute deviation from 0 of the time reversal symmetry metric of the optimized model responses. Interestingly, as we increased the number of layers, the models trained on the natural scenes and reversed contrast scenes tended to become more time reversal symmetric for the edges and sawtooth stimuli, while models trained on phase randomized and Gaussian scenes remained time reversal symmetric (**Fig. 6I, J, Supp. Fig. S6**). For *S*_14_, all models but the one trained on Gaussian scenes continued to break time reversal symmetry even as their depth increased (**Fig. 6K**). Thus, the more expressive models trained on natural scenes tended to break time reversal symmetries less than shallower models for the contrast asymmetric stimuli. Meanwhile, models trained on natural and phase randomized scenes broke time reversal symmetry for *S*_14_, while Gaussian scenes never resulted in trained models breaking time reversal symmetry.

To better understand these results, we also derived an analytical result for motion detectors that rely on different orders of correlations in the stimulus (**Appendix 2**). We found that spatially mirror symmetric detectors that rely on only pairwise correlations must be time reversal symmetric, while those relying on higher order correlations may break time reversal symmetry. Since motion detectors can use higher order correlations, if present, to best estimate motion (12), our analytical result suggests that the non-Gaussianity of the stimulus contrast is required (but not necessarily sufficient) for trained models to break time reversal symmetry with a given stimulus, in agreement with our numerical results (**Fig. 6**).

## Discussion

This research shows that fly motion detection algorithms break time reversal symmetry, that is, there are stimuli for which the response to the time reversed stimulus is not equal and opposite to the response to the original stimulus (**Figs. 1, 3, 4**). This property of the response does not seem to depend strongly on the symmetry features of the stimulus (**Fig. 4, Supp. Fig. S4**). However, machine learning and analytical models for motion detection each show that contrast asymmetries and higher-order correlations in training data can lead motion detection algorithms to break time reversal symmetry (**Figs. 5, 6, Appendix 2**). Interestingly, model complexity seems to suppress some types of time reversal symmetry breaking but not others (**Fig. 6**).

Time reversal is an under-studied symmetry in neuroscience. Some studies have used movies played in reverse to understand the encoding and different timescales of integration of human brain regions (33). Other studies have examined reversibility of dynamics in fMRI and in retinal recordings, where reversibility can be quantified in the degree to which state transitions obey detailed balance (34-37). Those studies found that the reversibility of neural dynamics can depend on stimulus and task, as well as on brain region. The framework presented here is distinct from prior work on reversibility in that it asks not about microscopic state transitions, but rather about the overall low-dimensional output of a well-studied neural computation. The symmetry breaking studied here requires the dynamical system to be driven and asks how its behavior changes under symmetry transformations of the driving stimulus. These properties appear distinct from microscopic reversibility. It would be interesting to pursue how algorithmic frameworks for neural computation relate to the state-space frameworks in terms of reversibility.

### Response time reversal symmetry breaking and light-dark asymmetrical processing

Both flies and humans process visual motion with algorithms that incorporate light-dark asymmetries (11, 38-40). The light-dark asymmetric processing has often been understood as reflecting optimization to natural scene statistics (12, 31, 32, 41). Is broken time reversal symmetry in motion responses simply another view of this contrast asymmetry in motion processing? The phenomena are related, but time reversal symmetry breaking is a broader category than the light-dark processing differences. Two lines of evidence point to a definitive distinction between the phenomena. First, the stimuli *S*_14_ and *S*_4_ are each contrast reversal symmetric but both result in responses that strongly break time reversal symmetry (**Figure 4**). These two examples demonstrate that time reversal symmetry breaking can occur even when all contrasts are balanced. Second, a large number of stimuli did not yield symmetrical responses under simultaneous contrast and time reversal (**Figure 4**). This double transformation of the stimulus reverses time and changes all edge types from light to dark and vice versa. The time reversal transforms light edges into dark ones, but then the contrast reversal transforms them back to light ones. Thus, responses that do not exactly invert under contrast-time reversal must also result from mechanisms that are distinct from mere differences in responses to light and dark moving edges. In sum, motion detectors with light-dark asymmetric responses must break time reversal symmetry, but time reversal symmetry breaking does not imply light-dark asymmetric responses. Thus, there is a broad set of stimuli that result in broken time reversal symmetry in responses, even when there is no light-dark asymmetry in the response (**Fig. 4, Fig. 6**).

### Mixing asymmetries in shallow networks

In interpreting machine learning models, we often expect symmetries in the training data to be reflected in the model. For instance, if training data for an orientation detector were isotropically distributed, we would expect an isotropic model, detecting all orientations equally. In fact, when that rotational symmetry is broken in a trained model, it is interpreted as reflecting the same broken symmetry in the stimulus (8). Interestingly, in the case we have studied here, time reversal symmetry in responses was *not* broken in our trained models by a broken time reversal symmetry in the training data — by construction, our training stimuli were time reversal symmetric (**Figure 5**). Instead, asymmetries and non-Gaussianity in the stimulus *contrast* distribution led to time reversal symmetry breaking. This mixing of contrast statistics with time reversal is clear in our analytical derivation (**Appendix 2**).

### Optimization and response symmetry breaking

Comparisons with optimized systems have proven useful in understanding the structure of motion detecting circuits and algorithms (42). A Bayes optimal motion detector will be sensitive to higher order correlations in the inputs over time and space (12), and our analytical result suggests that this sensitivity can cause a motion detector to break time reversal symmetry (**Appendix 2**). Our artificial neural networks use threshold linear rectifiers (**Fig. 5, 6**), which gives them access to both the second order correlations in canonical models and to higher order ones (31). Our results show that optimizing shallow neural networks to estimate velocity with natural scene inputs leads to time reversal symmetry breaking (**Figures 5, 6**), suggesting that optimization to natural scenes could explain the time reversal symmetry breaking in the fly.

Deeper, more expressive networks tended to reduce time-reversal symmetry breaking associated with contrast asymmetries (**Figure 6**). However, they did not reduce the response time reversal symmetry breaking of a stimulus that was light-dark symmetric. The deeper networks may be better able to extract the latent variable of velocity without considering the nuisance structure of the scene itself, but the specific symmetries retained or broken depend on the stimuli examined.

### The trained artificial neural networks do not reproduce fly percepts

We investigated how neural network architecture and training impact their time reversal symmetry to stimuli. Although the optimized networks taught us about the conditions that lead to breaking time reversal symmetry in responses, they produced responses that could or could not match the direction of the fly’s responses, depending on the stimulus parameters (**Supp. Fig. S5**). Prior work has shown that task optimization can recover features of the fly’s motion processing, especially when connectomic constraints are incorporated (30, 43). However, the tests here may depend on more subtle differences in processing than the comparisons made in prior work. The differences between artificial networks and the fly behavior could arise because our neural networks are missing response properties that constrain the fly’s responses: processing gain changes with luminance (44), temporal and spatial contrast adaptation (45, 46), and synaptic nonlinearities in motion detectors (29, 47-49). It seems *a priori* unlikely that a relatively narrow, shallow neural network using threshold-linear activation functions but trained on naturalistic data would produce net responses to specific visual stimuli similar to the full biological network’s behavioral output.

There are many structural (7), circuit (50), and computational (51, 52) parallels between flies and mammals in visual and motion processing. The artificial neural network studies here suggest that processing that breaks time reversal symmetry could arise naturally from an optimization process. It will therefore be interesting to examine patterns in how visual percepts break time reversal symmetry across phyla. The similar responses of flies and humans to peripheral drift illusions (17, 21), which break time reversal symmetry, suggest that other stimuli could also elicit similar responses.

Overall, this study has shown that biological motion detectors can break time reversal symmetry under a variety of conditions. In simulations, we showed that this broken symmetry can reflect constraints on the algorithm and its tuning to the input contrast distributions. Time reversal symmetry has been largely neglected in understanding sensory processing, but investigating when it is preserved or broken can help uncover the structure of sensory computations and how they are tuned to their inputs.

## Acknowledgments

We acknowledge helpful conversations with the Clark Lab, J. Demb, T. Emonet, J. Fitzgerald, C. Lynn, and J. Zavatone-Veth. This research was supported by NIH R01 EY026555 and NS121773.

## Contributions

NW and DAC conceived the new stimulus. NW obtained and analyzed behavioral data. MA obtained behavioral data. BZ performed neural network optimization and analysis. NW, BZ, and DAC wrote the paper.

**Supplementary Figure S1.**
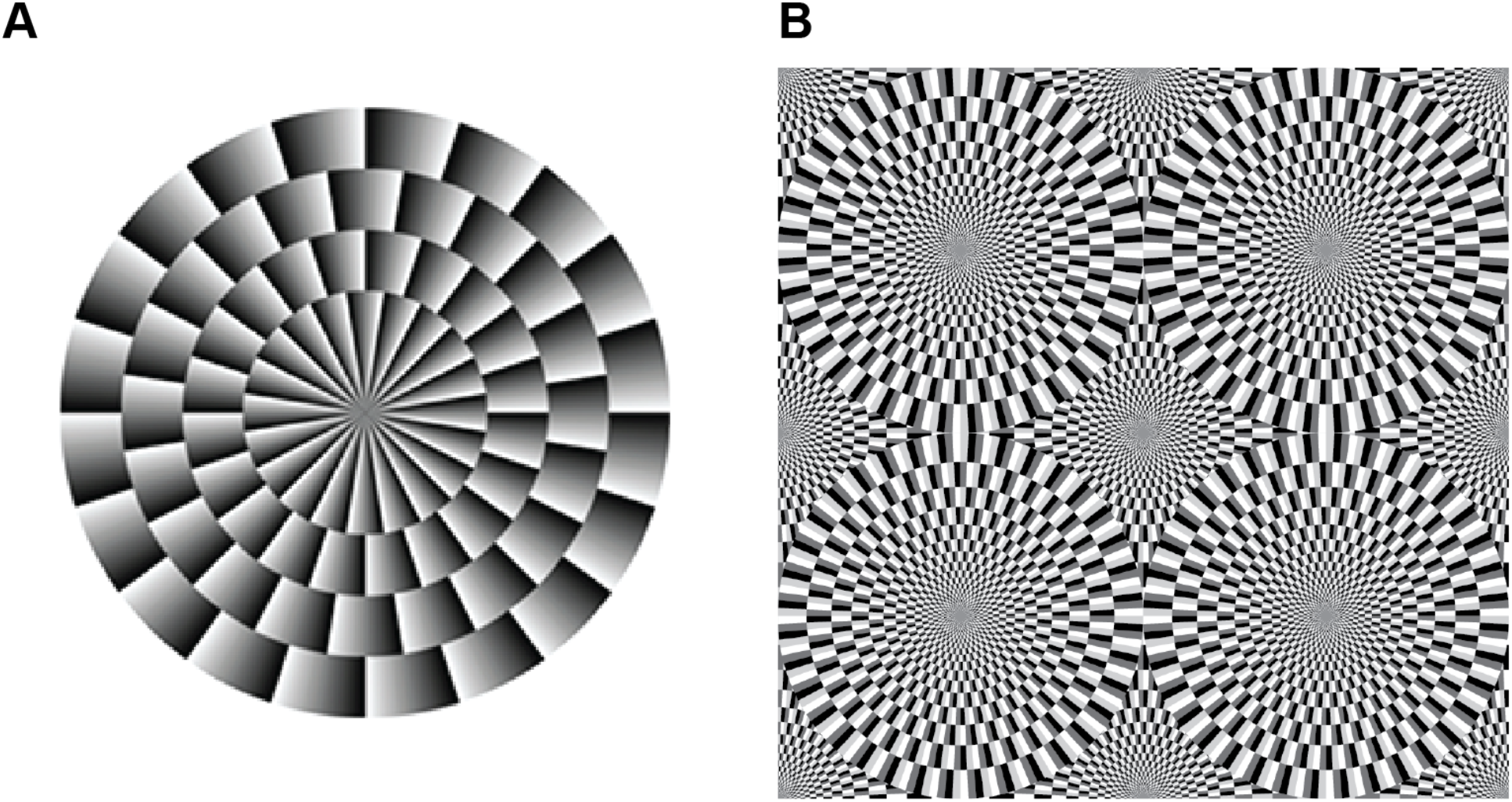
Stationary contrast stimuli that elicit motion percepts in humans. A. Peripheral drift illusion. When this stimulus is observed in the periphery of the visual field (i.e., not looking directly at it), it appears to rotate slowly clockwise. Adapted from (17). B. Snake illusion. When humans observe this stationary pattern, the wheels appear to rotate in different directions. This percept is enhanced when the eyes move. Adapted from (22).

**Supplementary Figure S2.**
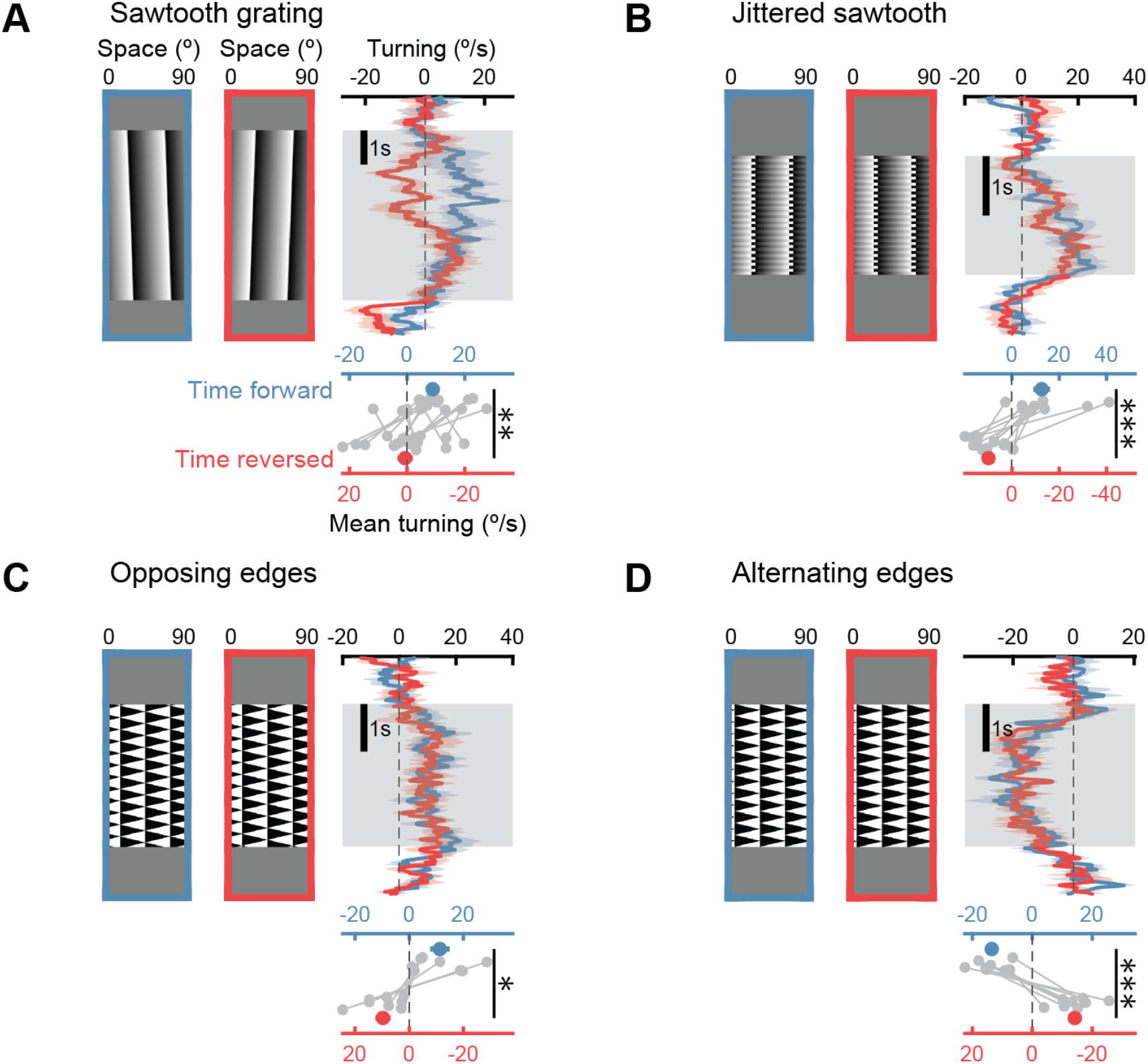
Additional stimuli that cause time reversal symmetry breaking in flies. A. As in **Figure 1D**, but for a version of the sawtooth stimulus drifting at 1.5 º/s. Time reversal symmetry breaking does not require that a stimulus be time-symmetric or stationary. N = 8 flies. B. As in **Figure 1D**, but for a jittered contrast sawtooth, which is time reversal symmetric. N = 8 flies. C. As in **Figure 1D**, but for an opposing edge stimulus (23), where light edges move in one direction and dark edges move in the other at the same points in time. The stimulus is the same when time is reversed. N = 8 flies. D. As in **Figure 1D**, but for an alternating edge stimulus, where light edges and dark edges alternate in time on the display, with light edges moving in one direction and dark edges in the other. This stimulus is very similar to the opposing edge stimulus (E), the only difference being that every other “column” of edges is phase shifted in time by half a period. However, it produces a very different response, with flies turning in the opposite direction from in (E). These two stimuli reveal that responses are not easily predicted by simply counting the net light and dark edges, since those are identical in these two stimuli. N = 8 flies. ^*^ P<0.05, ^**^ P<0.01, ^***^ P<0.001 by Student t-test.

**Supplementary Figure S3.**
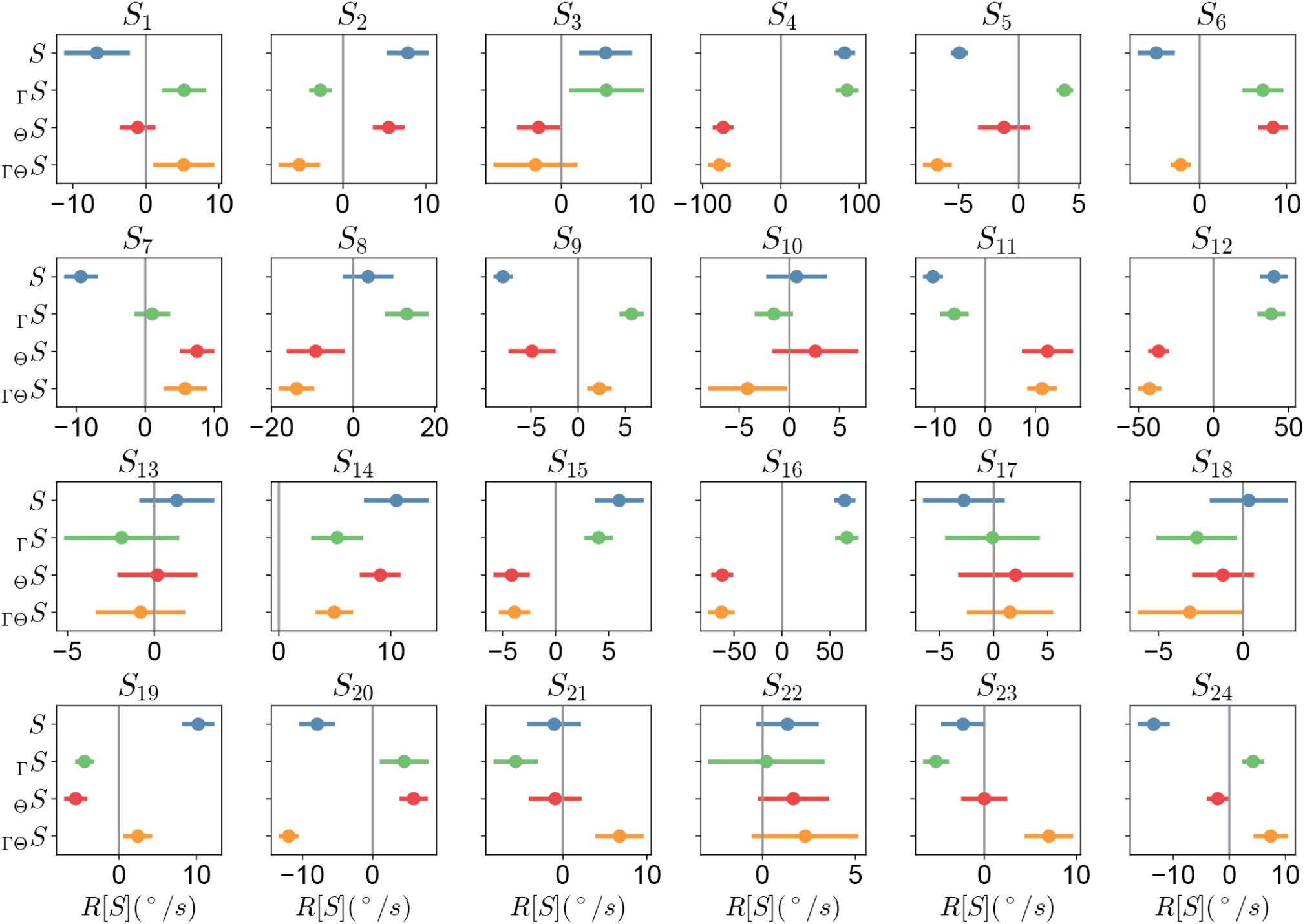
Mean responses ±SEM to all presented edge-tile stimuli. Edge-tile stimuli were presented for a total of three seconds, with each individual edge moving at 240°/s. Values here were averaged over the presentation period, then means and SEMs were computed over flies. *N* = 8-11 flies for each stimulus.

**Supplementary Figure S4.**
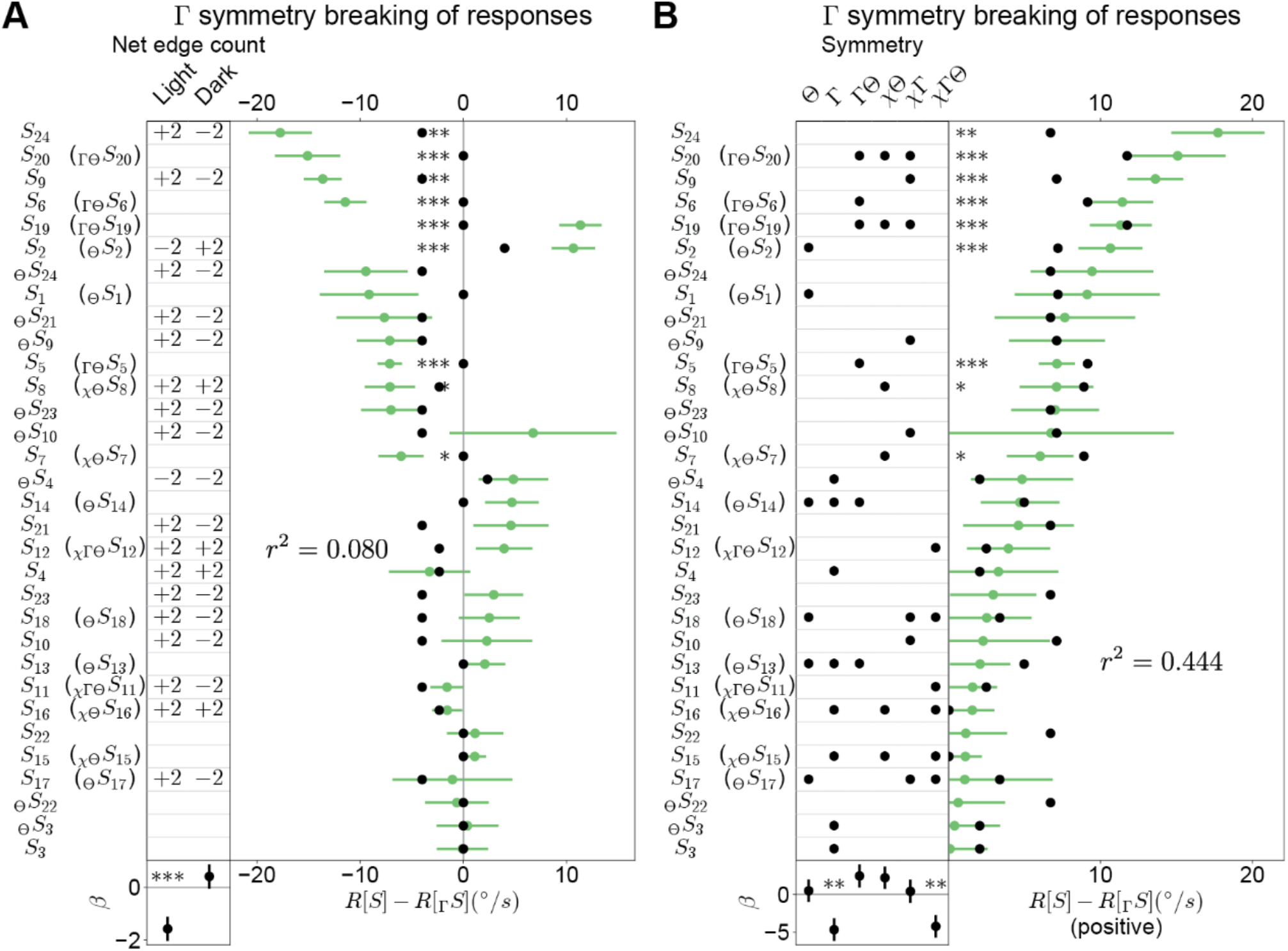
Contrast reversal symmetry breaking in the stimulus set. A. As in **Figure 4A**, but for contrast reversal symmetry breaking. Stimuli in parentheses indicate that the stimulus being tested possesses a symmetry (or multiple symmetries) that permit combining response asymmetry metrics from both *S* and 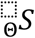 into a single measurement. B. As in **Figure 4C**, but for contrast reversal symmetry breaking.

**Supplementary Figure S5.**
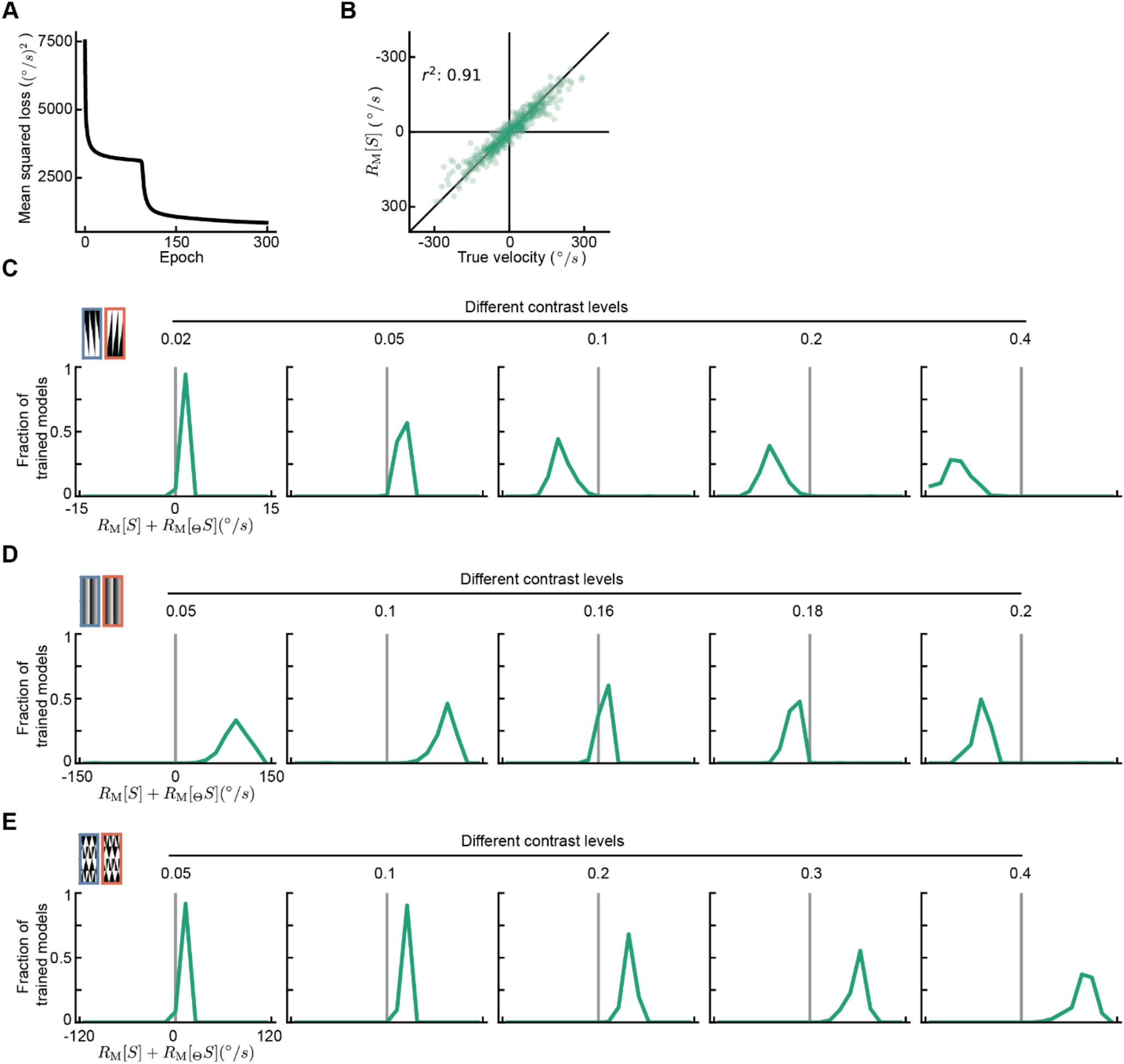
Direction of symmetry breaking depends on stimulus parameters. A. Example loss function over time during training. B. Performance of one of the trained models on the test data with a typical r-squared value of 0.91. Each dot represents a different test motion stimulus. C. As in **Figure 5J**, but for different contrast levels in the moving edge stimulus. D. As in **Figure 5K**, but for different contrast levels in the stationary sawtooth stimulus. E. As in **Figure 5L**, but for different contrast levels in *S*_14_.

**Supplementary Figure S6.**
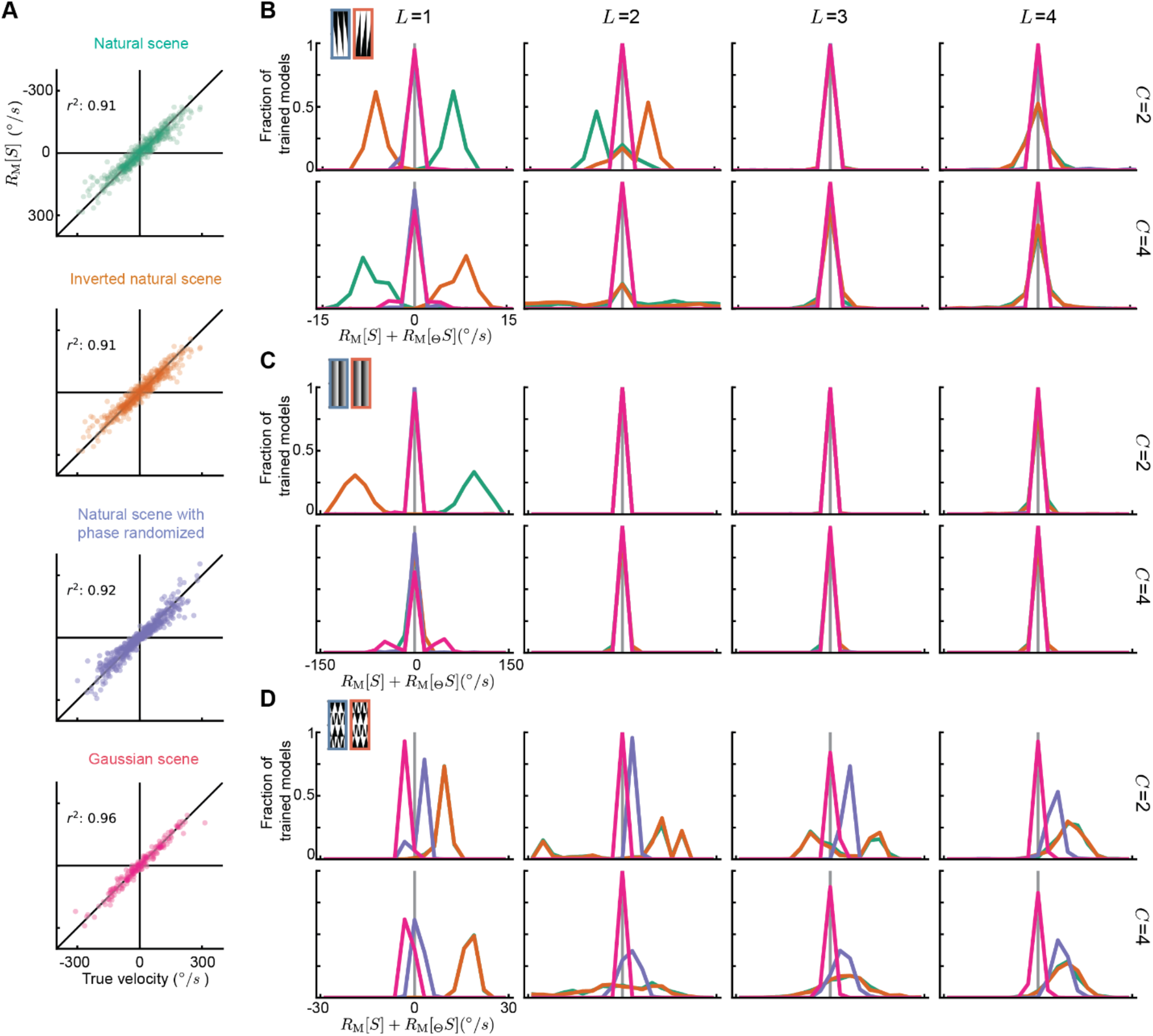
Raw model response distributions for different numbers of channels *C* and layers *L*. A. Typical plots of predicted vs true velocities for test data in single models trained on the four types of training data. B. Similar to **Figure 6B**, but for different numbers of channels *C* and layers *L*. C. Similar to **Figure 6C**, but for different numbers of channels *C* and layers *L*. D. Similar to **Figure 6D**, but for different numbers of channels *C* and layers *L*.

**Movie 1**. Movie illustrating the 24 edge-tile stimuli shown to the flies.

## Methods

### Data and code availability

All behavioral data in this paper is available along with code to make all the figures in this paper here: https://www.github.com/ClarkLabCode/TimeReversalSymmetry [to be made available on publication]. Code to run the machine learning portions of the analysis in this paper is also available in the same repository, as is code to generate and classify the edge tile stimuli. The behavioral data has also been deposited as a Dandi dataset available here: [to be made available on publication].

### Behavioral measurements

Wildtype *Drosophila melanogaster* vinegar flies (of strain Oregon R (53)) were grown at 20ºC, 50% relative humidity in a 12-hour-light, 12-hour-dark circadian cycle. Flies were collected on CO2 for behavior in their first 24 hours after eclosion, then tested 24 hours later, putting their age during testing at 36 to 48 hours. Their behavior was measured during the 3 hours after dawn or 3 hours before dusk.

To measure optomotor turning behavior, flies were tethered to a small needle that suspended them above an air-supported ball, which could rotate underneath them, as in prior experiments (54). Panoramic projection screens around them displayed green, monochrome stimuli (16) with a luminance of ∼100 cd/m^2^. Stimuli typically were presented in two 20-minute presentations of subsets of stimuli in random order.

### Visual stimuli

Visual stimuli were created in Matlab using Psychtoolbox (55-57). All stimuli were interleaved with periods of 2-3 seconds of mean gray luminance. In all cases, spatial phase of the stimulus was randomized for each presentation. The update rate for all stimuli was 180 frames per second.

Stimuli presented to flies in this research.

| Stimulus name | Description | Figures |
| --- | --- | --- |
| Drifting sinusoid | Sinusoidal contrast grating with wavelength of 30°, temporal frequency of 16 Hz, and 100% contrast. | 1D |
| Stationary spatial sawtooth | Stationary sawtooth grating in space with 45° period and 100% contrast. Gray interleaves. Originally published in (21). | 1E |
| Light edges | Periodic light edges moving in one direction. Spatial period of 30° with 100% contrast edges moving at 240°/s. | 1F |
| Uneven square wave grating | 100% contrast square wave grating with light and dark bars: 5° dark, 5° light, 25° dark, 25° light, so that there is an overall spatial period of 60°. Stimulus drifts at 30°/s. | 1G |
| Drifting spatial sawtooth | Sawtooth grating in space with 45° period and 100% contrast, drifting with a velocity of 1.5°/s, either in the direction of the | S2A |
|  | shallow gradient or in the direction of the steep gradient. |  |
| Jittered spatial sawtooth | Sawtooth grating in space with 45° period and 100% contrast. Jittered back and forth in space by 5° with a period of 1/10 second. | S2B |
| Opposing edges | Space-time pattern in which light edges move one direction and dark edges move the opposite direction simultaneously. Spatial period of 30° with 100% contrast edges moving at 240°/s. Originally published in (23). | S2C |
| Alternating edges | Space-time pattern in which light edges move one direction and dark edges move the opposite direction, with *only* light edges or dark edges occurring at any given time. Spatial period of 30° with 100% contrast edges moving at 240°/s. | S2D |
| Edge-tile stimuli | Periodic, 100% contrast stimuli created as described in **Methods** and in **Figure 2**. Each stimulus edge cell had a 30° extent and edges moved at 240°/s. | 3, 4 |

### Symmetry definitions

We define our stimuli, *S*(*x, t*), as the contrast at each point in space and time. There are three operators that reverse the time, space, and contrast of the stimulus: Θ, χ, and Γ. These operators act as follows:

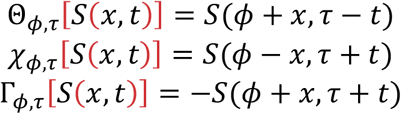

As a shorthand, we will typically drop the phase shift parameters and also denote the time reversed stimulus by a left subscript, so that,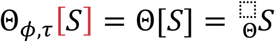. These phase shifts in space and time, *ϕ* and *τ*, may be chosen arbitrarily, and we will choose them specifically to identify symmetries in the stimuli.

A stimulus is defined as possessing a time reversal symmetry if Θ[*S*(*x, t*)] = *S*(*x, t*) for some pair of phase offsets *τ* and *ϕ*. Similarly, a stimulus possesses space reversal symmetry when χ[*S*(*x, t*)] = *S*(*x, t*) for some pair of offsets *τ* and *ϕ*, while a stimulus possesses contrast reversal symmetry when Γ[*S*(*x, t*)] = *S*(*x, t*) for some pair of offsets *τ* and *ϕ*. Each of these operators applied twice can result in a return to the original stimulus: 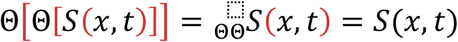 for pairs of offsets *τ* and *ϕ*. Similarly, 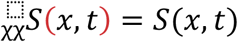 and 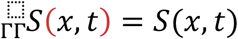 for pairs of offsets *τ* and *ϕ*.

As an example, a sinusoidal contrast grating drifting with a velocity *v* = *ω*/*k* is defined as *S*(*x, t*) = *c* sin (*kx* − *ωt*). This stimulus possesses contrast reversal symmetry because Γ[*S*(*x, t*)] = *S*(*x, t*) when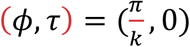. The stimulus also possesses space-time reversal symmetry, since 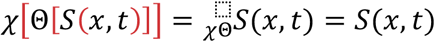 when 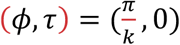 for the time reversal and (*ϕ, τ*) = (0,0) for the space reversal.

### Data analysis

Each fly’s responses were averaged over trials, and mirror symmetrized to generate a mean turning curve for each stimulus. Figure time traces show means and standard errors over flies. Time traces were averaged over the stimulus duration to create mean turning values for each fly and stimulus. The response, *R*, of a fly to each stimulus, *S*, is a scalar, *R*[*S*].

To assess symmetry breaking, we compared fly responses to each stimulus with their responses to the same stimulus transformed. Time reversal symmetry breaking was assessed by asking whether 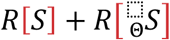 was different from 0. Contrast reversal symmetry breaking was assessed by asking whether 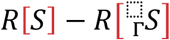 was different from 0. Time-contrast reversal symmetry breaking was assessed by asking whether 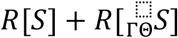 was different from 0. See below for definitions of the symmetry transformations. Statistical significance of the symmetry breaking was computed using a Student t-test, which was Holm-Bonferroni corrected for the 6 different symmetries assessed for each stimulus.

### Making a suite of stimuli with specific spatial, temporal, and contrast symmetries

We set out to create a suite of periodic stimuli possessing different combinations of symmetries, as defined above. Each stimulus in this novel suite of stimuli, collectively called edge-tile stimuli, is characterized by a *m* by *n* rectangular arrangement of moving edges, with *m* moving edges across the temporal dimension and *n* moving edges in the spatial dimension. Each moving edge “tile” within the *m* by *n* arrangement is one of four types: a rightwards light edge, a leftwards light edge, a rightwards dark edge, or a leftwards dark edge (**Fig. 2A**). The entire stimulus consists of many *m* by *n* unit cells that are tessellated across both space and time.

Every stimulus with a unit cell of size *m* by *n* is uniquely defined by the tuple (*m, n*) (the stimulus’s shape) and a vector *v*_2_ of length *m* × *n* × 2, with each element in the vector being either 1 or −1. Every individual moving edge tile within the unit cell is encoded by two elements within the vector, with the first element of the pair encoding the polarity of the edge (1 for a light edge and −1 for a dark edge) and the second encoding the direction of the edge (1 for rightwards and −1 for leftwards). The first two elements in *v*_2_ encode the leftmost edge at the earliest time in the unit cell. Each successive pair of elements correspond to edges as they go from left to right, then from first-in-time to last-in-time.

Reversal transformations (time reversal, space reversal, contrast reversal) to each stimulus and their associated operators (Θ, χ, and Γ, respectively) can be operationalized as matrix multiplication. Let the notation *v*_2_ represent the vector that defines stimulus *S*, for any *S*. For each shape (*m, n*) and each operator *O*[*S*] on stimulus *S*, one may compute a square matrix *A*_*o, m, n*_ such that *A*_*o, m, n*_ × *v*_2_ = *v*_*o* [2]_, for all *S* of shape (*m, n*). Given *O, m*, and *n*, the matrix *A*_*o, m, n*_ can be procedurally generated.

Because stimuli unit cells are tessellated infinitely in space and time (**Fig. 2B, C**), translating the pattern could result in a new ordering of edges (and what would appear to be a new unit cell) without changing the true, tessellated stimulus. Thus, different encoding vectors could define the same stimulus. To address this issue, translations in space and time can also be operationalized as matrix multiplication, just as with reversal transformations. Every stimulus of shape (*m, n*) has *m* × *n* different encoding vectors that are all equivalent to one another, and consequently, *m* × *n* unique square matrices *B* where *B* × *v*_2_ is a vector that encodes an equivalent stimulus to *v*_2_ (including the identity matrix). We call this set of matrices the tessellation matrix set. The tessellation matrix set for a given shape can be generated by first finding the matrices *B* _*ϕ, m, n*_ and *B*_*τ, m, n*_, where *B*_*ϕ, m, n*_ × *v*_2_ encodes *S* translated by one cell in space and *B*_*τ, m, n*_ × *v*_2_ encodes *S* translated by one cell in time. The tessellation matrix set *T* is defined as follows:

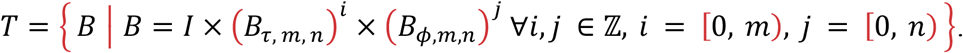

There are 2^*m*×*n*×2^ = 4^*m*×*n*^ different possible encoding vectors for unit cells of shape *m* × *n*. A stimulus corresponding to encoding vector *v*_2_ is *O* symmetric (where *O* is an operator) if there exists a matrix *B* in the *m* × *n*-sized tessellation matrix set where *B* × *A*_*o,m,n*_ × *v*_2_ = *v*_2_. To create a suite of stimuli with specific spatial, temporal, and contrast symmetries, all stimuli of shape (4, 2) and of shape (2, 4) were created and categorized according to whether they were Θ, Γ, χ, ΓΘ, χΘ, χΓ, and χΓΘ symmetric. 24 different stimuli were selected from all generated stimuli, excluding those that were χsymmetric: Four patterns that exhibited no symmetries at all, and two patterns that exhibited every other unique combination of symmetries (**Table 1**, *bottom*). The patterns were chosen to minimize discontinuities in time (for instance a dark region that instantaneously becomes light). When possible, stimuli for each combination of symmetries were selected to have both non-zero and zero net motion over time and space.

To give concreteness to these definitions of *v*_;_ and the symmetry operators *A*_Θ_, *A*_χ_, and *A*_Γ_, below we give specific examples of these for the case of a (2,4) (time, space) shaped unit cell. In the vector *v*_;_, the edge tile polarity *p*_*ij*_ and direction *d*_*ij*_ are each ±1.

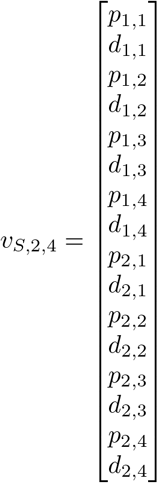

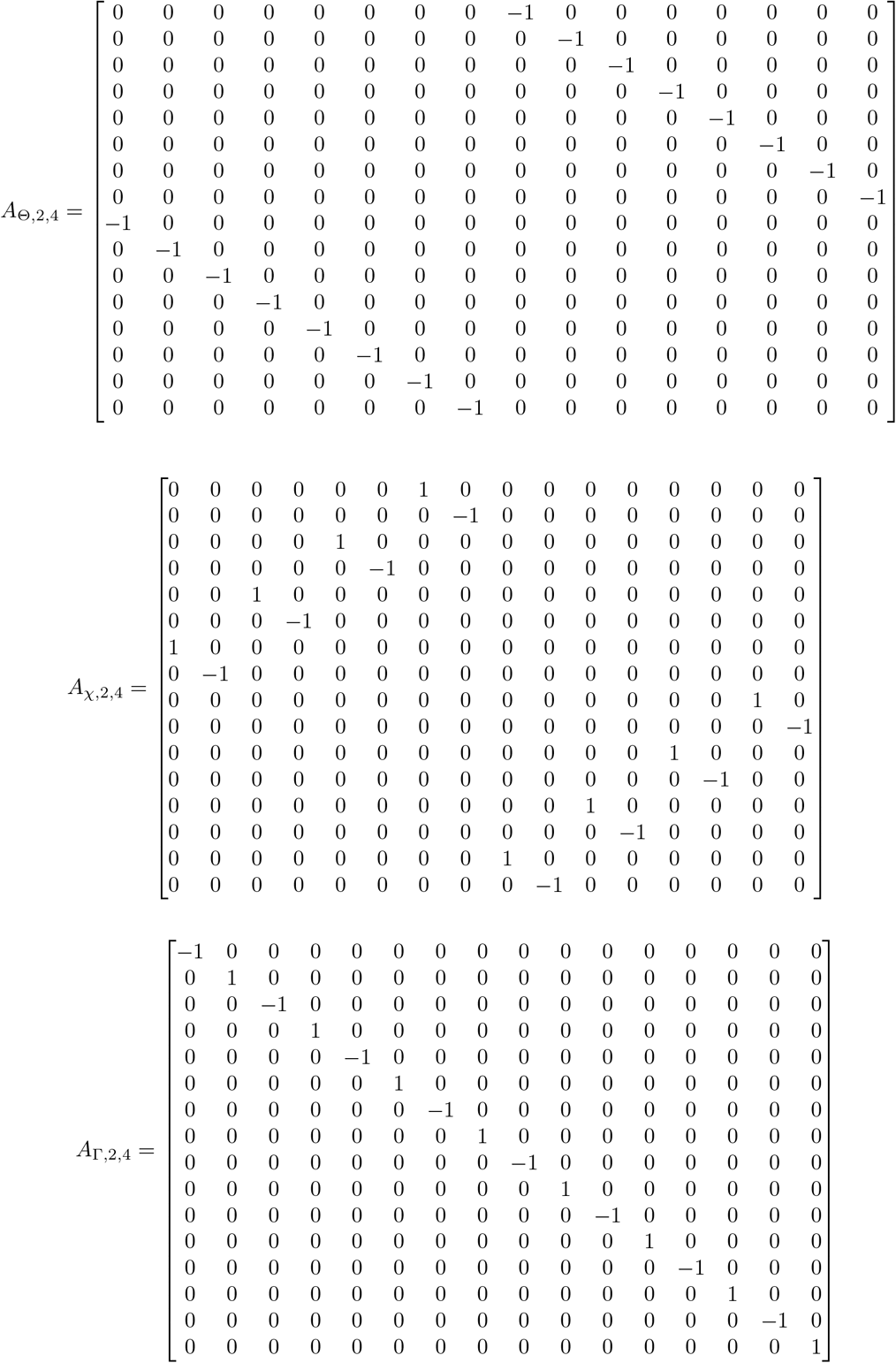

### Training data for artificial neural networks

Natural scene data was adapted from an online database with a total of 241 images (27). Each image in the dataset is a 251 × 927 grey scale matrix and spans 360º panoramically in azimuth and 97.5º in elevation. One ommatidium of the fly compound eye has an acceptance angle of roughly 5º, and thus, each image was filtered by a 2-dimensional Gaussian kernel with a full width at half maximum of 5º. Within each image, contrast was calculated by subtracting the local mean and divided the local mean. The local mean was obtained by using a 2-dimensional Gaussian kernel with the standard deviation of 30º.

In this study, only yaw motion was considered, and we simulated the images moving with a velocity trajectory over time. The velocity trajectory was generated by an Ornstein-Uhlenbeck process with an autocorrelation half-decay time of 0.2 s (autocorrelation time constant of 0.2/ln(2) s) and a standard deviation of 100º/s. Each generated velocity trajectory has a length of 200 steps with a time resolution of 0.01s, from which only the last 49 steps of the velocity traces were used to generate synthetic moving images. For each image, 100 different velocity trajectories were generated, and the image was shifted horizontally according to these velocity trajectories. Thus, there were 100 movies for each image and each movie was 0.5s long (50 steps with a time resolution of 0.01s). Every frame of the movies was subsampled to a matrix with size 20 × 72, and each pixel spanned roughly 5º in both azimuth and in elevation. Since only horizontal motion was considered, for each movie, single row slices were selected at random. In the end, one sample of data had a shape of 50 × 1 × 72 (50 time points by 1 by 72 spatial locations with 5º spacing). For each movie, there was a counterpart with time reversed and the spatial coordinate reversed. In total, there were 144000 training samples and 48800 testing samples.

For **Figure 6**, the contrast reversed natural scene was generated simply by multiplying the natural scene samples by –1. The phase randomized natural scene samples were created by Fourier transforming the original natural scene samples, setting the magnitude corresponding to the zero frequency to zero, adding random phases to other modes, and inverse Fourier transforming back. The standard deviation of this distribution was set to match the standard deviation of the pixel values in the natural scene contrasts. This process preserves the second order spatial correlations in the scene, since the power spectrum is preserved. The phase randomized scenes are not Gaussian. When a non-Gaussian image moves, it generates higher order spatiotemporal correlations not determined by the second-order correlations alone. The Gaussian samples were generated by sampling pixel values independently from a Gaussian distribution. The standard deviation of this distribution was set to match the standard deviation of the pixel values in the natural scene contrasts. When a Gaussian image moves, higher order spatiotemporal correlations provide no information beyond that in the second-order correlations, since the first and second moments of a Gaussian distribution determine all higher moments. In the case of the phase randomized and Gaussian images, the corresponding movies were generated exactly as in the natural scene case, but no contrast was calculated, since all image values were already centered at 0.

### Evaluation stimuli for artificial neural networks

We used three stimuli to evaluate the time reversal symmetries of model responses: moving edges, stationary sawtooth contrast patterns, and *S*_14_. For the moving edge stimuli, each movie contained 12 equally spaced light edges moving from left to right (forward) or right to left (backward). The velocities of the moving edges were set to be 60º/s. The positive contrast of the moving edges were set to 0.05 and the negative contrast set to –0.05 in the main figures before being smoothed by a Gaussian kernel with a full width at half maximum of 5º (as we filtered the images). We varied the contrast scales to have the values of 0.02, 0.05, 0.1, 0.2, 0.4 (**Supp. Fig. S5**). For the static sawtooth stimuli, the period of the sawtooth pattern was set to be 45º and the maximum/minimum contrasts were 0.05/–0.05 in the main figure before being smoothed by a Gaussian kernel with a full width at half maximum of 5º. We varied the contrast scales to have the values of 0.05, 0.1, 0.16, 0.18, 0.2 (**Supp. Fig. S5**). For *S*_14_, we used the same method as in the section *Visual Stimulus*, and set the maximum/minimum contrasts to be -0.05/0.05 in the main figure before spatial filtering. We also varied the contrast scales to have the values of 0.05, 0.1, 0.2, 0.3, 0.4 (**Supp. Fig. S5**).

### Model architecture of artificial neural networks

Convolutional neural network models were built to predict the velocities for the input movies generated according to the last section. The number of channels *C* and number of layers *L* were varied: *C* = 2, 4 and *L* = 1, 2, 3, 4. The channels of the first layer had a depth of 50, and its overall shape was 50 × 1 × 3 (time extent by vertical extent by horizontal extent). The channels of the subsequent layers (if any) had an overall shape *C* × 1 × 3. In these cases, the three spatial inputs represent the three spatial inputs to the motion detectors in the fly (28), as in prior task optimization in the fly visual system (30). For each channel, there is a counterpart channel with the last dimension flipped to enforce the opponency, where the output of the counterpart channel was subtracted from the original channel. The output layer of each model was a spatially uniform average of the last convolutional layer. To enforce the left-right symmetry in space, the final output of the model contains two parts. The first part was the predicted velocity of the original sample, and the second part was the predicted velocity of the sample flipped horizontally (the last dimension of the sample). The final output was simply the difference between the first part and the second part. In this way, if a stimulus was flipped horizontally in space, the velocity predicted by the model was enforced to equal to the original response multiplied by -1.

### Artificial neural network model training and testing

The weights in each model architecture were trained until convergence with 100 different initializations of the parameters. Coefficient of determination, or *r*^2^, was reported for the test dataset. We then asked how the model responded to the evaluation stimuli. Response time reversal symmetry breaking was assessed by plotting the time reversal symmetry metric, 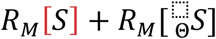, for the trained models with *r*^2^ larger than 0.8 for each architecture and training data type. This threshold ensures that we only examine models for which the training worked to a significant degree.

## Appendix 1: Time reversal symmetries in canonical models to detect motion

In the examples below, we will consider the space-time averaged responses of different classical models to a spatiotemporal stimulus *S*(*x, t*).

### The Hassenstein-Reichardt Correlator (HRC) model (1)

In the HRC model for motion detection, signals from two adjacent points in space are split, one signal is delayed relative to the other, and then the delayed and non-delayed signals from the two points in space are multiplied, before the products are subtracted. The net result is a signal that computes the difference in spatiotemporal correlations in two directions in space-time. When this response is averaged over space and time, it can be represented alternatively in Fourier space as:

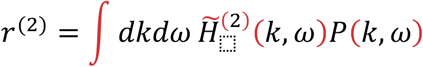

where *P*(*k, ω*) is the power spectral density of the stimulus in spatial and temporal frequency space and 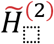 is a weighting function. (See the definitions in **Appendix 2**.) The weighting function 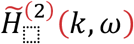 is mirror-symmetric in space because of the subtraction in the model, so that 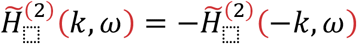. When time is reversed, it’s the equivalent of inverting *ω* in the stimulus. Therefore, the space-time averaged response 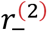 to a time reversed stimulus is (see the derivation also in **Appendix 2**.):

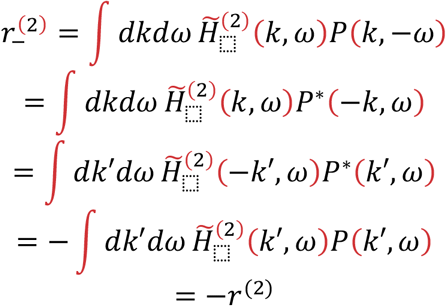

Therefore, in the HRC model, reversing time leads to an equal and opposite mean response.

### *The motion energy model* (2)

This model is very similar to the HRC above, in that it is sensitive only to pairwise correlations in the stimulus. Like the HRC, mean responses are simply a weighting function of the power spectrum of the stimulus. Also like the HRC, the motion energy model is spatially mirror-symmetric, so that the derivation above also applies directly to it. Therefore, motion energy models likewise invert their space-time averaged responses when time is reversed.

### *Optimized gradient model* (3)

Prior work has asked how an optimal model would estimate velocity with Gaussian input statistics and Gaussian noise. That work found that under low signal-to-noise conditions, the dominant term in the optimal estimator senses only pairwise correlations in the stimulus. But under high signal-to-noise conditions and rigid body image translation, they found that the optimal estimate of the true image velocity is:

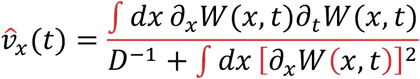

Where *W*(*x, t*) is a functional acting on the stimulus, *S*(*x, t*). When time is reversed, only the temporal derivative of *W* is inverted, and all other terms remain the same. Since the numerator in this expression is proportional to the (spatially integrated) temporal derivative of *W*, playing the stimulus backward inverts the motion estimate of this gradient-detector model.

### *Lucas-Kanade optic flow computation* (4)

The Lucas-Kanade model is a classical model for estimating optic flow, based on computing the displacement that minimizes the error between a movie frame and a displaced, subsequent frame. For rigid body translation of an image in one dimension, this calculation supposes that

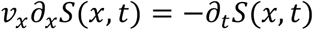

An estimate for *v*_*x*_ may be computed by solving for the best fit value of *v*_*x*_ over all space. The best fit solution is:

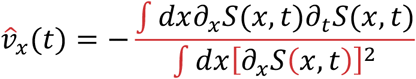

Here, as in the optimal solution from Potters and Bialek (3), the solution is proportional to the temporal derivative of the image intensity. When a movie is played in reverse, that temporal derivative is inverted, and all velocity estimates are likewise inverted. Thus, this algorithm for computing optic flow also obeys time reversal symmetry.

## Appendix 2: Only sensitivity to correlations beyond second-order can confer time reversal symmetry breaking

Motion detectors work by being sensitive to different spatiotemporal correlations in the stimulus. Suppose there is a stimulus *S*(*x, t*), defined over space and time. The motion energy model and the Hassenstein-Reichardt correlator (HRC) model (1, 2) are examples of models that are sensitive to only second order spatiotemporal correlations. In that case, we have two spatiotemporal filters, *H*_1_ and *H*_2_, that act on the stimulus to create filtered signals *A*_1_ and *A*_2_:

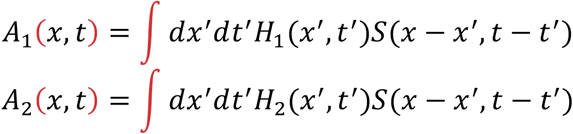

(In the motion energy model, *H*_1_ = *H*_2_.) The mean response over time and space of a multiplier unit in the HRC model (or a squared input unit in the motion energy model) is:

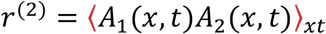

Where the mean is taken over *x* and *t*: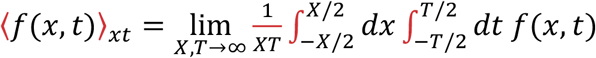, where *X* is the spatial extent and *T* is the temporal extent of the integration.

When we calculate these convolutions instead in Fourier space, we find:

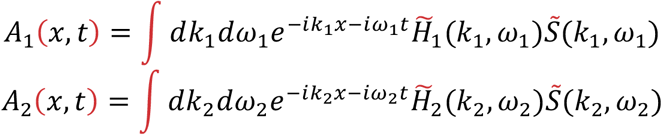

We substitute the above two equations into the equation for *r*^(2)^, and integrate over space and time to find

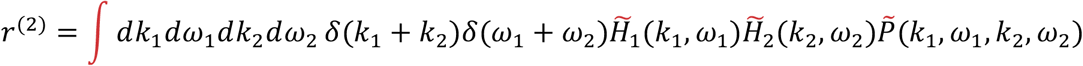

where *δ*(·) is the Dirac delta function and 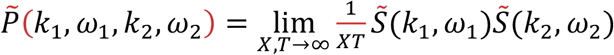 is the power spectral density of the signal (58, 59). We perform one integral over spatial and temporal frequencies to find:

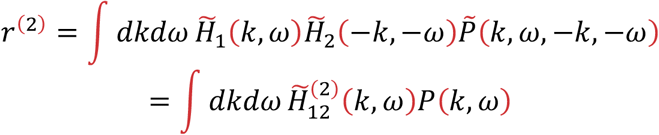

where we define 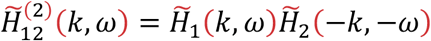 and the power spectral density of the stimulus becomes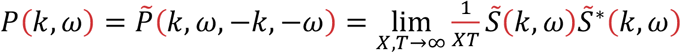. We use the fact that 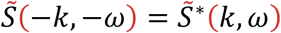 for Fourier transforms of real-values functions.

Now, more generally, one can define 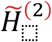 for the sum or difference of some set of HRC or motion energy units, so that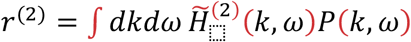. In these models, mirror-symmetric units are subtracted to create spatial mirror-symmetry. Here, we suppose that the world is left-right symmetric and that the left-right mirror-symmetry is imposed on our models. To flip the weighting function 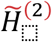 in space, we simply invert its spatial frequency argument.

Then, the spatial mirror-symmetry of the model is represented as:

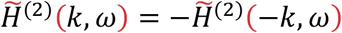

Now, we examine how this model responds if the stimulus is reversed in time, that is, that we present the model with 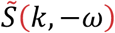 instead of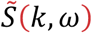. The space-time averaged response to the temporally reversed stimulus, 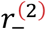, is:

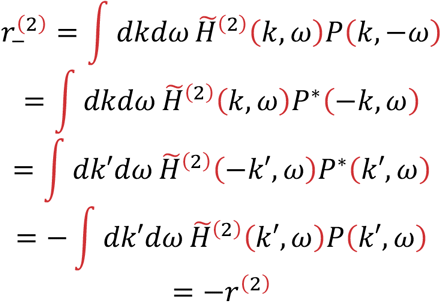

In the third line, we substituted *k′* = −*k* and adjusted the limits of integration appropriately so that it still integrates from −∞ to ∞. The equality above indicates that the mean response to the stimulus is reversed when the stimulus is reversed in time. Fundamentally, this equality relies on the spatial anti-symmetry of the weighting function and on the fact that *P* = *P*^*^.

We can generalize this to models that are sensitive to correlations between triplets of points over space and time. In that case, the mean response to a stimulus is:

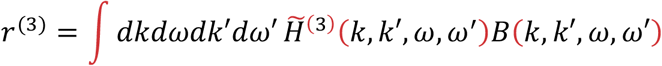

where we define 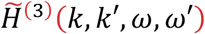 as a weighting function over frequency pairs and the bispectrum density (60) is defined as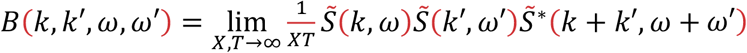. We assume the same left-right anti-symmetry: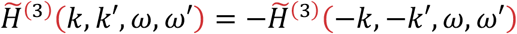. If we reverse the time of the stimulus, using the same logic as the prior equation, we find:

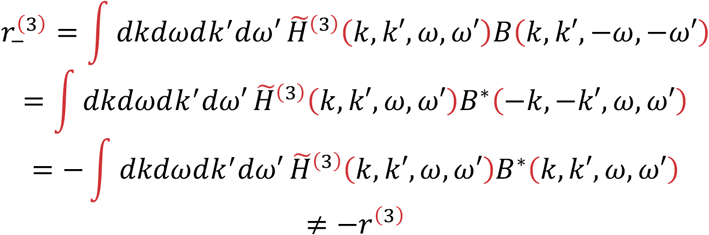

The inequality arises because in general *B* ≠ *B*^*^.

Similar arguments can be applied to further higher-order correlations. For instance, the mean response to fourth-order products of the stimulus is a weighted average of the trispectrum (60). For the bispectrum, the trispectrum, and beyond, complex conjugation does not in general yield the same function. This means that the pairwise correlation case is special in generally obeying time reversal symmetry. Motion detectors sensitive to any correlations beyond second-order need not invert their responses on time reversal, depending on the structure of the stimulus. But in the case of a motion detector sensitive to only pairwise correlations, it *must* obey time-reversal symmetry for all stimuli.

